# Phosphorylation of Z granule components regulates germ granule organization and RNAi inheritance

**DOI:** 10.64898/2026.10.06.756944

**Authors:** Lizaveta Pshanichnaya, Martin Brehm, Ivan Mikicic, Svenja Hellmann, Aaquib Mohd, Petra Beli, Sebastian Falk, René F. Ketting

## Abstract

Small RNAs represent important carriers of epigenetic information in nematodes, and factors involved in their biogenesis reside within perinuclear germ granules. Among these condensates, Z granules are essential for the inheritance of RNA interference (RNAi). The mechanisms by which Z granules form and promote this process remain unclear. We show that Casein kinase I alpha (KIN-19) is present in Z granules and associated with DEPS-1 and HERD-1 phosphorylation. In the absence of DEPS-1 and HERD-1 phosphorylation, animals have enlarged Z granules and enhanced RNAi inheritance. We propose that DEPS-1 acts as a phosphorylation-regulated scaffold that bridges ZNFX-1 and HERD-1 proteins. Our study demonstrates that Casein kinase I alpha-mediated phosphorylation is an important factor in germ granule modulation and RNAi inheritance, and additionally provides a novel molecular scaffold for understanding germ granule complexity in *C. elegans*.

## Introduction

Epigenetic marks, such as DNA methylation, histone methylation and small non-coding RNA, alter gene expression in a way that is heritable in the absence of changes in DNA sequence(Perez and Lehner, 2019; Sarkies, 2020; Grishok, 2021; Fitz-James and Cavalli, 2022). Due to inheritance of these marks, stimuli such as viral infection, starvation, or environmental stress experienced by the parent can affect progeny that never encountered the original trigger(Rechavi *et al*., 2014; Klosin *et al*., 2017; Lowe, Shukla and Kennedy, 2025).

In the nematode *C. elegans,* silencing of messenger RNAs (mRNAs) by double-stranded RNA, a process known as RNA interference (RNAi), can persist for multiple generations(Fire *et al*., 1998; Grishok, Tabara and Mello, 2000). During the initial response, the endonuclease RDE-8 cleaves the target mRNA and MUT-2 (also known as RDE-3) adds a 3′ poly(UG) tail, promoting the recruitment of RNA-dependent RNA polymerases (RdRPs), which synthesize abundant secondary 22G RNAs(Sijen *et al*., 2001; Tsai *et al*., 2015; Shukla *et al*., 2020). Subsequently, these small RNAs are loaded onto worm-specific Argonautes (WAGOs) and transmitted via egg and sperm to propagate RNAi to the next generation(Schreier *et al*., 2022). A key step in inheritance is the re-establishment of silencing in the embryo (Schreier *et al*., 2025). One of the main factors in this process is a protein ZNFX-1, which contains a central UPF1-like superfamily 1 helicase domain and a chain of six zinc finger motifs (Ishidate *et al*., 2018; Wan *et al*., 2018; Schreier *et al*., 2025). ZNFX-1 orthologs are present across diverse animal species, including humans, where it is involved in innate immunity(Cheng and Parker, 2025). In *C. elegans*, ZNFX-1 associates with transcripts that have acquired poly(UG) tails and directs their localization to germ granules (Wan *et al*., 2018; Ouyang, Zhang and Seydoux, 2022). In the inheriting generation, ZNFX-1 is required to sustain poly(UG)ylation and sRNA amplification(Wan *et al*., 2018; Ouyang, Zhang and Seydoux, 2022), and in its absence, targeted genes fail to be silenced(Schreier *et al*., 2025).

Maintenance of silencing, along with other steps of small RNA biogenesis, takes place within germ granules, phase-separated organelles that assemble through liquid–liquid phase separation and remain closely associated with the germ cell nucleus(Phillips and Updike, 2022; Uebel, Rajeev and Phillips, 2023). Proper spatiotemporal regulation of small RNA biogenesis is believed to be achieved through the compartmentalization of distinct factors into specific types of germ granules. For instance, proteins localized to Mutator foci primarily drive small RNA production, whereas those within P granules contribute to gene repression(Marnik and Updike, 2019; Uebel *et al*., 2020; Scholl, Liu and Seydoux, 2024). ZNFX-1, in turn, is enriched in Z granules, which are specifically implicated in the inheritance of small RNA-driven responses(Wan *et al*., 2018), and is required for the re-establishment of silencing in RNAi-inheriting offspring (Schreier *et al*. 2025). Recent studies using TurboID proximity-labeling expanded our understanding of the protein composition of these granules (Zhao *et al*., 2024; Li *et al*., 2025; Chen *et al*., 2026). However, gaps remain in our understanding of how these proteins interact with one another, how these interactions are regulated and how these factors regulate small RNA production and function.

Besides the RNA-binding proteins ZNFX-1 and WAGO-4, Z granules contain proteins that do not bind RNA directly but instead regulate protein-protein interactions to ensure proper timing and localization of small RNA biogenesis(Wan *et al*., 2018). HERD-1, for instance, is a Z granule component that has been proposed to prevent mixing of germ granules(Zhao *et al*., 2024). In *herd-1^-/-^* mutant, P granules, Z granules and Mutator foci components have been described to merge, accompanied by persistence of RNAi inheritance for over 50 generations, compared with 3-5 generations in wild-type animals. The protein DEPS-1, which is also present in Z granules(Huang *et al*., 2025), directly interacts with *C. elegans* PIWI protein PRG-1 (Suen *et al*., 2020). This interaction is important for proper PRG-1 function but is not required for PRG-1 granular localization(Suen *et al*., 2020). A yeast two-hybrid screen identified an interaction between DEPS-1 and EDG-1. Reducing EDG-1 expression by RNAi enlarges DEPS-1 foci without affecting PRG-1. We note, and will describe in the results, that EDG-1 bears strong sequence similarity to HERD-1. Another structural component, PID-2 (also known as ZSP-1), localizes to the periphery of Z granules(Placentino *et al*., 2021; Wan *et al*., 2021). In *pid-2^-/-^* mutant animals, Z granules become enlarged and less numerous. Two redundant eTudor-domain proteins, PID-4 and PID-5, were discovered to coimmunoprecipitate with PID-2. *pid-4^-/-^;pid-5^-/-^* double mutants were found to phenocopy *pid-2^-/-^* mutants, suggesting that PID-2 functions mostly through PID-4 and PID-5 or *vice versa*(Placentino *et al*., 2021). Beyond the identity of all these Z granule-residing proteins, not much is known about how they mutually affect each other and how they affect Z granules and RNAi inheritance.

A range of protein-protein interactions drives granule formation and regulates the reactions that occur within them, and post-translational modifications such as phosphorylation can modulate these contacts(Li *et al*., 2022). In *C. elegans* embryos, phosphorylation of MEG proteins plays an important role in regulating P granule condensation and dissolution(Wang *et al*., 2014). Phosphorylation can regulate protein-protein interactions directly or through mediators such as 14-3-3 proteins(Stevers *et al*., 2018; Huang *et al*., 2022). This family of phospho-binding proteins can promote or disrupt interactions to control liquid-liquid phase separation. Although no such interactions have yet been reported in *C. elegans*, the human RNA-binding protein SMAUG1 interacts with 14-3-3 proteins via four phospho-regulated short linear motifs, and these interactions promote the dissolution of SMAUG1 condensates(Fehilly *et al*., 2023). The role of phosphorylation in multi-condensate systems, such as the highly complex germ granules in *C. elegans*, has not been well-studied.

Here, we characterize the Casein kinase Iα KIN-19 and the phosphopeptide-binding adaptor 14-3-3 protein PAR-5 as components of germ granules: KIN-19 is found within Z granules, whereas PAR-5 is localized in foci close to but not overlapping with them. Granular localization of both proteins depends on PID-4 and PID-5. Our analyses show that animals in which KIN-19 localization to Z granules is compromised exhibit reduced phosphorylation of the Z granule components HERD-1 and DEPS-1. Mutants lacking these phosphorylation sites form enlarged Z granules and have enhanced RNAi inheritance. Moreover, we propose a model in which DEPS-1 bridges ZNFX-1 and HERD-1 within a single complex. Consistent with this model, loss of DEPS-1 abolishes the granular localization of HERD-1 and reduces ZNFX-1 granular intensity. Disruption of DEPS-1 and ZNFX-1 interaction leads to ZNFX-1 destabilization. These studies begin to reveal how specific interactions and their regulation affect germ granule properties and the RNAi inheritance process that occurs within them.

## Results

### KIN-19 and PAR-5 localize to distinct germ granule compartments

We previously identified the eTudor domain proteins PID-4 and PID-5 as partially redundant components of *C. elegans* Z granules(Placentino *et al*., 2021). Deletion of both genes, but not of either single mutant, disrupted Z granule morphology and 22G RNA homeostasis. To further explore the biological functions of PID-4 and PID-5, we re-examined immunoprecipitation (IP) data from our previous study. In both datasets, we identified two highly conserved proteins, the casein kinase Iα ortholog, KIN-19, and the 14-3-3 protein PAR-5 that binds phosphorylated motifs to modulate protein function (Fig. 1A,B). Both proteins are crucial for embryogenesis: KIN-19 as a component of the Wnt signaling pathway(Walston *et al*., 2004; Banerjee *et al*., 2010) and PAR-5 as a factor necessary for cell polarity(Mikl and Cowan, 2014; Wu, Espiritu and Rose, 2016). Our findings suggested that a phosphorylation-dependent mechanism may operate within Z granules, something that has not been previously addressed. To explore this, we tagged KIN-19 and PAR-5 with GFP. The N-terminal region of PAR-5 mediates homodimerization, and our attempt to tag PAR-5 C-terminally led to animal sterility, indicating that modifications at either terminus disrupt PAR-5 function. Accordingly, we inserted the GFP sequence internally within the disordered loop between two alpha-helices; AlphaFold3 predictions suggest that GFP remains flexible in this arrangement (Fig. EV1A,B)(Obsil and Obsilova, 2011). Confocal microscopy revealed that KIN-19 is found in the germline cytoplasm and is also enriched within germ granules, with additional signal detected in the spermatheca (Fig. 1C,EV1C). Based on the morphology of the signal, KIN-19 may be present on the apical membrane of this structure, which is known to be highly folded (Wormatlas). PAR-5 displayed a broader expression pattern, with signal also present in the nucleus and with slight enrichment at the cell membranes within the germline (Fig. 1D, EV1D).

**Figure 1.**
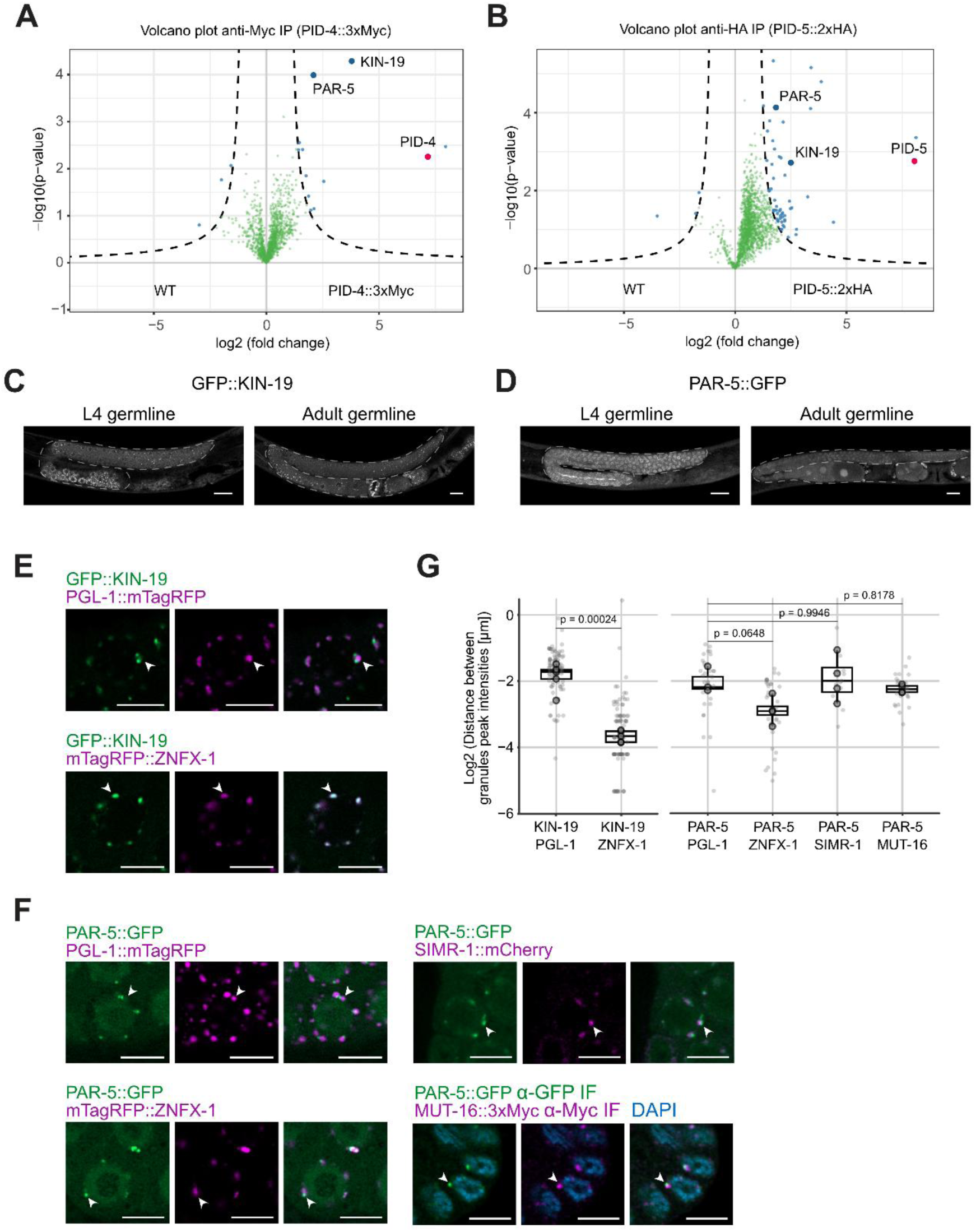
Granular localization of KIN-19 and PAR-5. (A-B) Adopted data from the previous publication (Placentino *et al*., 2021). Volcano plot showing proteins enriched in PID-4::3xMyc (A) and PID-5::2xHA (B) immunoprecipitations (IP), quantified using label-free proteomics. Dashed lines indicate thresholds for significance (p = 0.05) and two-fold change in enrichment, curvature of the enrichment threshold c = 0.05. Dots indicate proteins that are enriched and depleted (blue), or detected (green), bait protein is indicated in red. **(C-D)** Representative fluorescence microscopy of GFP::KIN-19 (C) and PAR-5::GFP (D) in L4 and Adult animals. Dashed lines represent the germline boundary. Images are representative of three independent biological replicates. Scale bars, 20 μm. **(E-F)** Representative fluorescence microscopy of the distal germline region in young adult hermaphrodites expressing GFP::KIN-19 together with PGL-1::mTagRFP or mTagRFP::ZNFX-1 (E) and PAR-5::GFP together with PGL-1::mTagRFP, mTagRFP::ZNFX-1, SIMR-1::mCherry or MUT-16::3xMyc (immunostaining)(F). Arrows indicate individual perinuclear germ granules. Images are representative of three independent biological replicates. Scale bars, 5 μm. **(G)** Quantification of distances between peak intensities of granules shown in (E-F). Each light dot represents the distance between peak intensities of individual granules; dark dots represent the mean distance between granules within one worm; the box represents the 25th percentile and 75th percentile of the per-worm means, and horizontal bars indicate median values of the per-worm means, whiskers reach the most extreme non–outlier values. For logarithmic data representation, 0 values were replaced with half the pixel size value. Statistical significance was assessed using Welch’s t-test for KIN-19 comparisons and Dunnett’s multiple comparison test for PAR-5 comparisons. n = 3-5 biological replicates.

To define the specific granules containing KIN-19 and PAR-5, we generated strains co-expressing established germ granule markers and imaged them by enhanced-resolution confocal microscopy (Zeiss Airyscan). KIN-19 signal visually appeared to overlap strongly with Z granule marker ZNFX-1 and showed significantly shorter distances to ZNFX-1 than to the P granule marker PGL-1, indicating that KIN-19 resides within Z granules (Fig. 1E,G). In contrast, PAR-5-containing granules were separated from ZNFX-1 granules and also distinct from PGL-1, SIMR-1, and MUT-16, suggesting that PAR-5 localizes to untested D or E granules, or a novel germ granule compartment (Fig. 1F,G). Taken together, these results demonstrate that the kinase KIN-19 and the 14-3-3 protein PAR-5 are present in distinct germ granules, even though they were both identified as PID-4/PID-5 interacting proteins(Placentino *et al*., 2021).

### KIN-19 and PAR-5 granular localization depends on PID-4 and PID-5

To investigate the functional relationship between KIN-19 and PAR-5, and PID-4 and PID-5, we examined how loss of PID-4 and PID-5 affected the granular localization of KIN-19 and PAR-5. Given the known redundancy of PID-4 and PID-5, we focused on the *pid-4^-/-^;pid-5^-/-^* double mutant(Placentino *et al*., 2021). Confocal imaging of the pachytene region in young adult germlines revealed that the number of KIN-19-positive granules, normalized to the number of P granules, was significantly reduced in the *pid-4^-/-^;pid-5^-/-^* double mutant (Fig. 2A,C). KIN-19 did retain association with Z granules in the *pid-4^-/-^;pid-5^-/-^* mutant, but its granular intensity was markedly reduced, while ZNFX-1 intensity in these granules was increased (Fig. 2B,D,EV2A,B). Interestingly, the granular localization of PAR-5, that is distinct from Z granules, was also significantly reduced (Fig. 2E,F). We therefore conclude that PID-4 and PID-5 are required for the efficient recruitment of KIN-19 and PAR-5 to germ granules.

**Figure 2.**
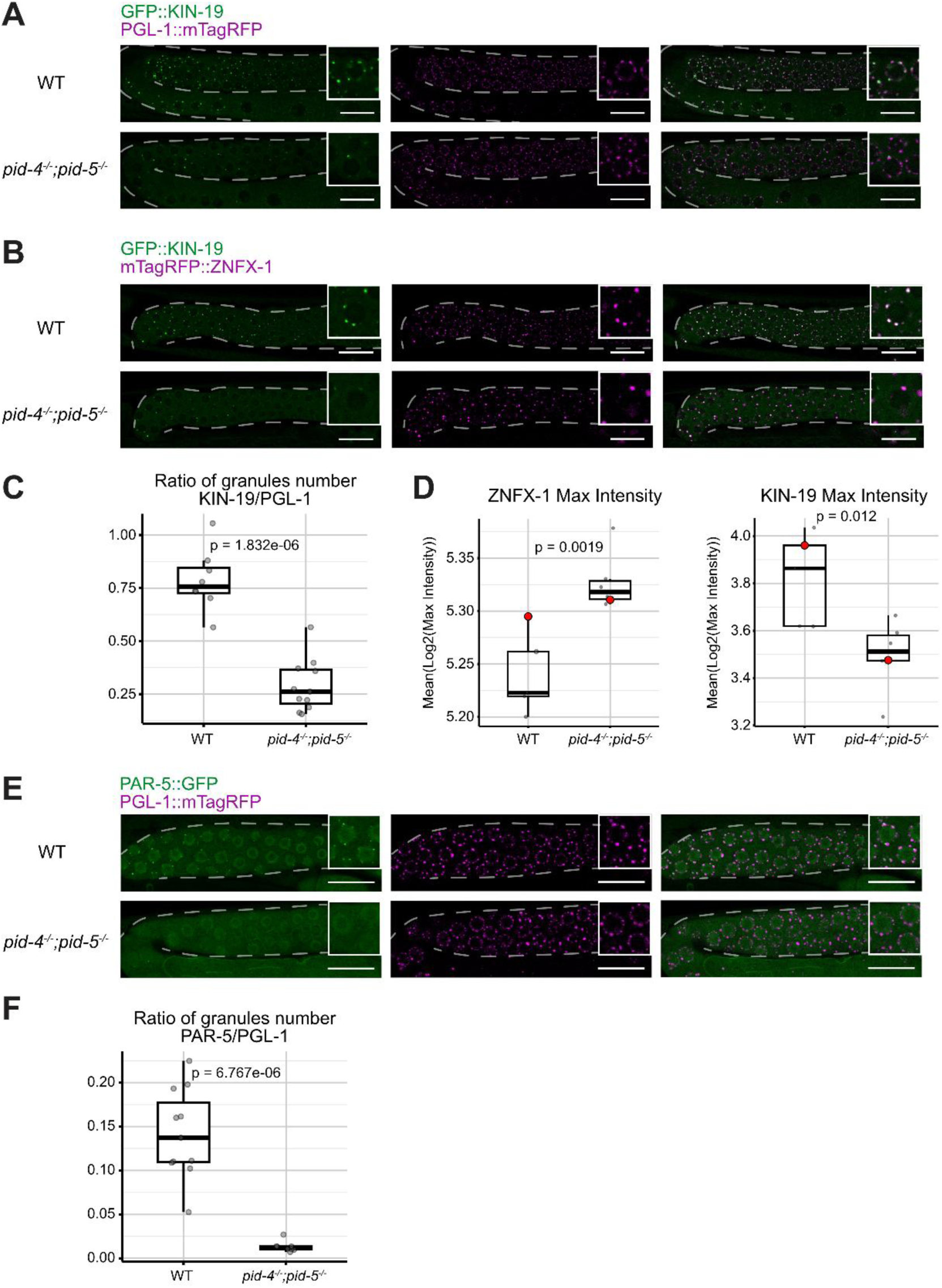
The number of KIN-19 and PAR-5 granules is significantly reduced in *pid-4^-/-^;pid-5^-/-^* double mutant. (A-B,E) Representative fluorescence microscopy of GFP::KIN-19 (A-B) and PAR-5::GFP (E) together with PGL-1::mTagRFP (A,E) or mTagRFP::ZNFX-1 (B) in young adult animals in the indicated mutant backgrounds. Dashed lines represent the germline boundary. n = 8-11 biological replicates. Scale bars, 20 μm. **(C,F)** Quantification of granule number in the pachytene region of young adult germ line shown in (A,E). The number of KIN-19 (C) and PAR-5 (F) granules was normalized to the number of PGL-1 granules. Each dot represents the granule ratio for an individual animal; boxes indicate the 25th–75th percentile range, whiskers reach the most extreme non–outlier values, and horizontal lines indicate the median. Statistical significance was assessed using Welch’s t-test. n = 8-11 biological replicates. **(D)** Quantification of microscopy images shown in (B). Dots represent the mean of log2-transformed maximum granule intensity per worm, boxes indicate the 25th–75th percentile range, and horizontal lines indicate the median, whiskers reach the most extreme non–outlier values. Red dots refer to representative micrographs shown in (B). Statistical significance was assessed using a linear mixed model. n = 5-6 biological replicates.

### DEPS-1 and HERD-1 are hypo-phosphorylated in *pid-4^-/-^; pid-5^-/-^* double mutants

According to previous studies, both KIN-19 and PAR-5 are essential for embryonic development(Banerjee *et al*., 2010; Mikl and Cowan, 2014). Additionally, the broad expression of both proteins suggested their involvement in other cellular processes besides a potential role in RNAi. Thus, disruption of KIN-19 and PAR-5 expression generally could affect many pathways and complicate the identification of their function within germ granules. However, the reduction of KIN-19 and PAR-5 presence in granules in the *pid-4^-/-^;pid-5^-/-^* double mutant provided a unique opportunity to dissociate their granule-specific functions from their broader cellular roles.

To identify granule-associated phosphorylation events, we performed phosphoproteome analysis on wild-type and *pid-4^-/-^;pid-5^-/-^* double mutant animals. As KIN-19 localized in Z granules, we examined whether the phosphorylation levels of Z granule components identified in the ZNFX-1 TurboID proximity labeling experiment were altered (Zhao *et al*., 2024). We found that phosphorylation in regions containing multiple phosphorylated residues of DEPS-1 (including T606 and S608) and HERD-1 (including S712 and S713) was strongly and significantly reduced in the mutant background (Fig. 3A,B). DEPS-1 and HERD-1 protein levels were not significantly changed in the *pid-4^-/-^;pid-5^-/-^* mutant (Fig. 3C). Despite the high protein sequence similarity of HERD-1 and EDG-1, that we address later, only HERD-1 phosphorylation was reduced. Residue S713 is conserved between HERD-1 and EDG-1, but according to our dataset, it is not phosphorylated in EDG-1. We conclude that in *pid-4^-/-^;pid-5^-/-^* double mutant animals, phosphorylation of DEPS-1 (most likely residues T606 and S608) and HERD-1 (most likely residues S712 and S713) is reduced.

**Figure 3.**
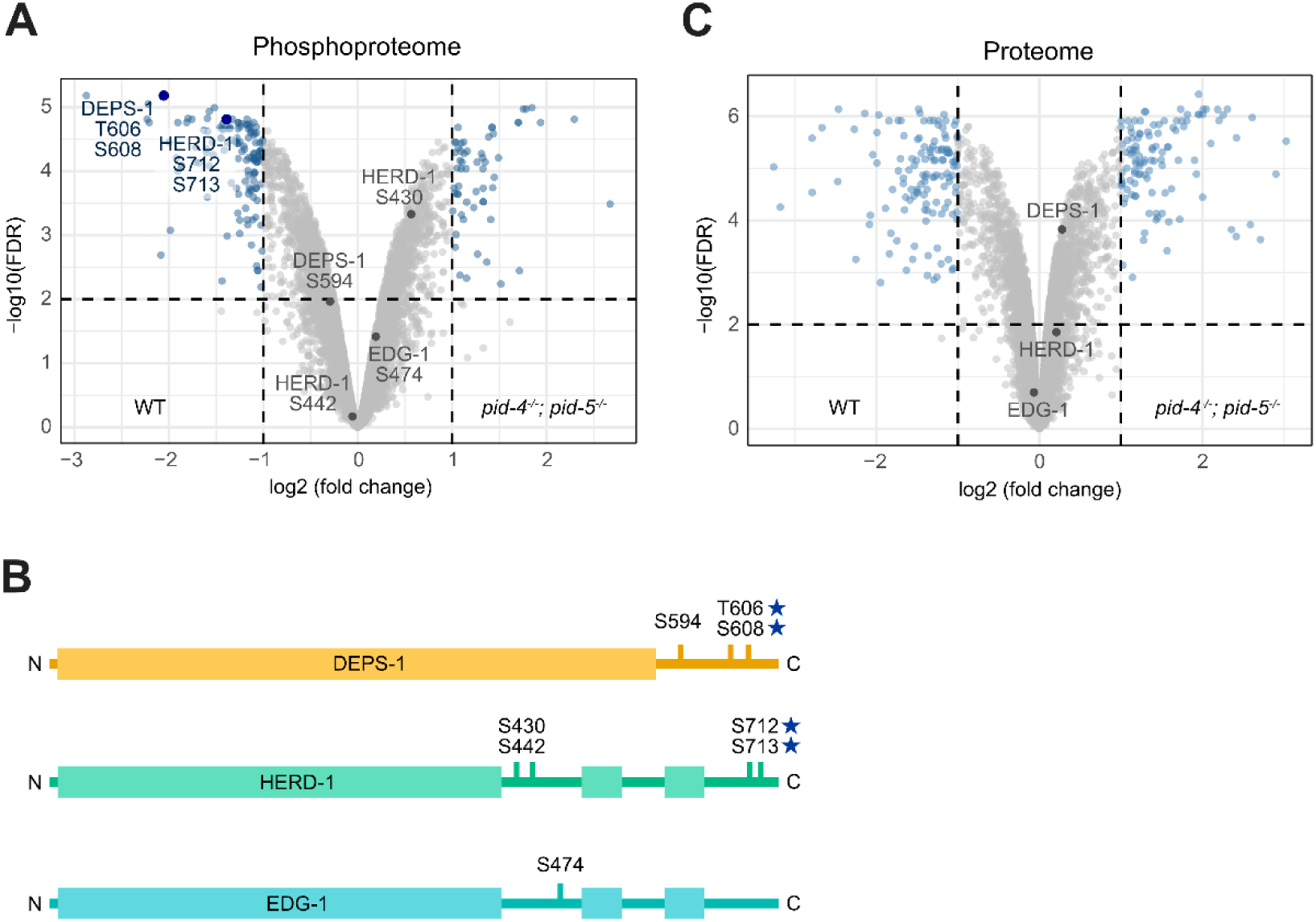
The reduction of KIN-19 and PAR-5 presence in granules leads to changes in DEPS-1 and HERD-1 phosphorylation levels. (A,C) Volcano plot showing differential phosphorylation sites (A) or protein (B) abundance in *pid-4^-/-^;pid-5^-/-^* animals versus wild-type. Experiments were performed on native protein extracts from young adult hermaphrodites (n = 5 biological replicates). The x-axis shows the mean fold change in abundance for each protein in *pid-4^-/-^;pid-5^-/-^* samples relative to wild-type. The y-axis represents the −log₁₀(FDR) of observed enrichments. Dashed lines indicate thresholds for significance: FDR = 0.01 and a two-fold change in enrichment. Dots indicate proteins that are enriched or depleted (blue), or detected (grey). **(B)** Scheme of DEPS-1, HERD-1 and EDG-1 proteins highlighting phosphorylation sites. Structured domains are shown in thick lines and disordered regions in thin lines. Stars mark phosphorylation sites that are less phosphorylated in *pid-4^-/-^;pid-5^-/-^* double mutant.

### ZNFX-1 granules are affected by DEPS-1 and HERD-1 phosphorylation

The DEPS-1 and HERD-1 phospho-sites that are affected in the *pid-4^-/-^;pid-5^-/-^* double mutant are present in the C-terminal disordered regions of these proteins. We mutated them to study the role of DEPS-1 and HERD-1 phosphorylation specifically, separated from other potential substrates of KIN-19. First, we removed the possibility of phosphorylation by mutating relevant serine and threonine residues to alanine: DEPS-1 (T606A; S608A), referred to as DEPS-1 phospho-dead, and HERD-1 (S712A; S713A), referred to as HERD-1 phospho-dead. Second, we generated phospho-mimic mutants in which the relevant serine and threonine residues were substituted with aspartate, whose negative charge mimics (but is not equal to) the phosphorylated state: DEPS-1 (T606D; S608D), referred to as DEPS-1 phospho-mimic, and HERD-1 (S712D; S713D), referred to as HERD-1 phospho-mimic.

We then measured the mean intensity of germ granules in mutant animals. As ZNFX-1 granules show increased intensity closer to the germline turn, this positional gradient could bias intensity measurements (Fig. 4A). We therefore analyzed the mid and late pachytene regions of the germline separately, as Z granule intensity was more consistent within each region. These regions were selected at the same distance from the germline turn across all images. Granule number was counted across the entire pachytene region, since this measure is less sensitive to differences in granule intensity.

**Figure 4.**
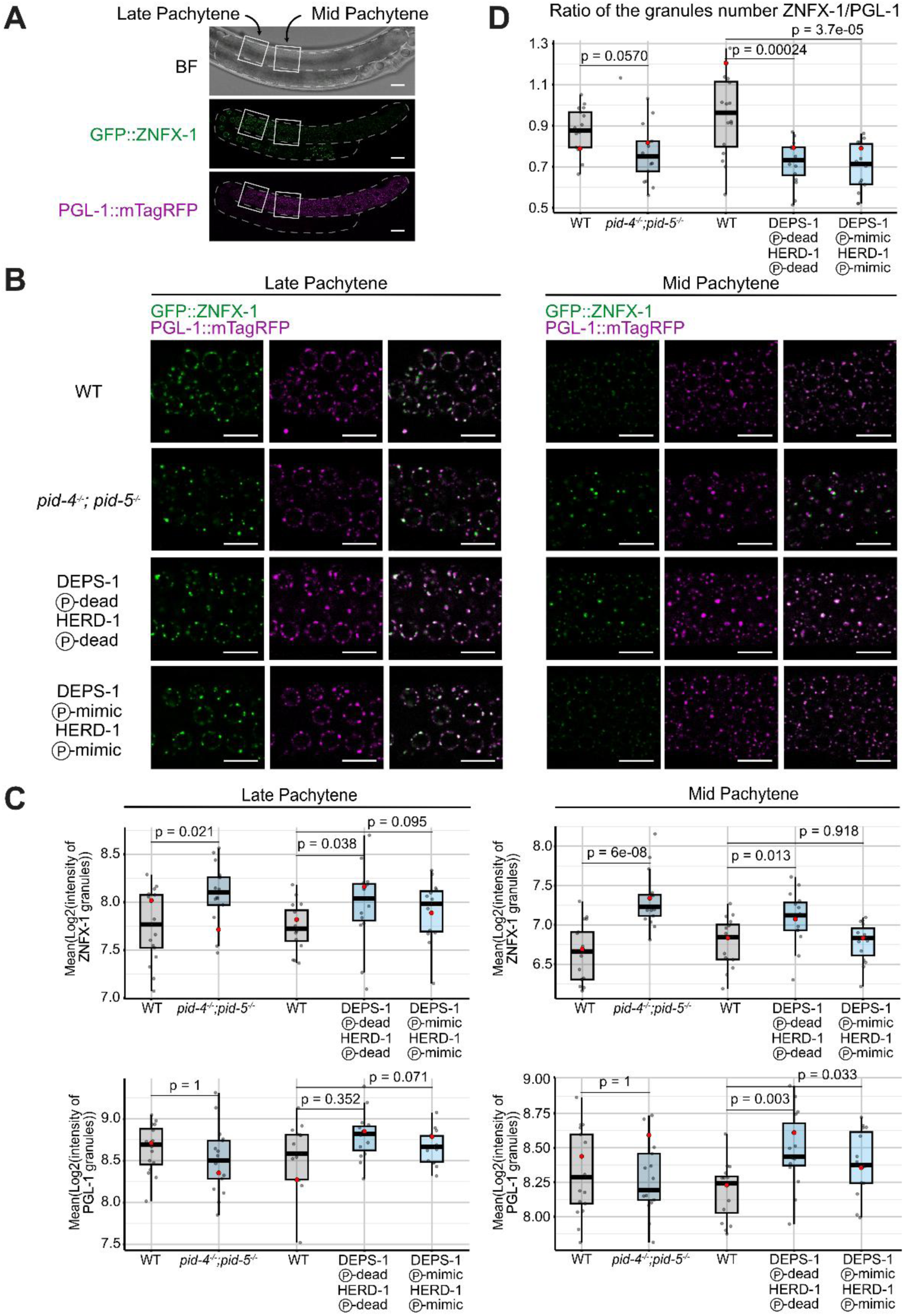
Z granule intensity depends on DEPS-1 and HERD-1 phosphorylation. **(A)** Representative fluorescence microscopy of GFP::ZNFX-1 and PGL-1::mTagRFP in adult animals. Dashed lines represent the germline boundary, and squares indicate Mid and Late Pachytene regions of the germline. Images are representative of three independent biological replicates. Scale bars, 20 μm. **(B)** Representative fluorescence microscopy of GFP::ZNFX-1 and PGL-1::mTagRFP in Mid and Late Pachytene regions of the same young adult animals in the indicated backgrounds. DEPS-1 (P)-dead corresponds to DEPS-1 (T606A;S608A) and HERD-1 (P)- dead to HERD-1 (S712A;S713A). DEPS-1 (T606D;S608D) is DEPS-1 (P)-mimic and HERD-1 (S712D;S713D) is HERD-1 (P)-mimic. Scale bars, 10 μm. **(C)** Quantification of ZNFX-1 and PGL-1 granule intensity shown in (B), analyzed separately for the Mid and Late Pachytene regions. Each dot represents the mean log2-transformed granule intensity of an individual animal; boxes indicate the 25th–75th percentile range, and horizontal lines indicate the median, whiskers reach the most extreme non–outlier values. Red dots refer to representative micrographs shown in (B). Statistical significance was assessed using a linear mixed model, with Holm’s correction for multiple comparisons when needed. n = 15–17 animals per condition. **(D)** Quantification of granule number in the whole pachytene region of the germline. The number of ZNFX-1 granules was normalized to the number of PGL-1 granules. Each dot represents the granule ratio for an individual animal; boxes indicate the 25th–75th percentile range, and horizontal lines indicate the median, whiskers reach the most extreme non–outlier values. Statistical significance was assessed using Welch’s t-test for *the pid-4^-/-^; pid-5^-/-^* comparison with wild-type and Dunnett’s multiple comparison test for the comparison of DEPS-1, HERD-1 phosphorylation mutants with wild-type. n = 15–17 biological replicates.

According to our previous work, and confirmed in this manuscript, ZNFX-1 granule intensity increased in *pid-4^-/-^;pid-5^-/-^* mutants in both mid and late pachytene regions, while PGL-1 was unaffected (Fig. 4B,C)(Placentino *et al*., 2021). In DEPS-1 and HERD-1 single phosphorylation mutants, we did not observe changes in the mean intensity of ZNFX-1 and PGL-1 granules (Fig. EV3A-B). However, ZNFX-1 granule intensity did increase in the mid and late pachytene regions when we combined DEPS-1 phospho-dead and HERD-1 phospho-dead mutants, similar to our observations in *pid-4^-/-^;pid-5^-/-^* mutants (Fig. 4B-C). The phospho-mimic mutants did not show this effect (Fig. 4B-C). These results suggest that a major cause of the Z granule changes in *pid-4^-/-^;pid-5^-/-^* double mutants is DEPS-1 and HERD-1 hypo-phosphorylation.

In some cases, phospho-dead and phospho-mimic mutants produced similar effects, suggesting that these effects are not driven directly by phosphorylation status. For instance, in DEPS-1;HERD-1 phospho-dead and phospho-mimic double mutants, unlike *pid-4^-/-^;pid-5^-/-^* mutant, PGL-1 granules have higher intensity in the mid pachytene region (Fig. 4B-C). In addition, in DEPS-1 single phospho-dead and phospho-mimic mutants, the ratio of ZNFX-1 and PGL-1 granule numbers in the entire pachytene region was reduced (Fig. EV3C). The same effect was observed for DEPS-1; HERD-1 phospho-dead and phospho-mimic double mutants (Fig. 4D). A potential explanation for these effects could be altered expression of DEPS-1 and HERD-1 in phospho-dead and phospho-mimic mutants. Consistent with this hypothesis, we observed that HERD-1 protein level was reduced in phospho-dead and phospho-mimic mutants (Fig. EV3D). These blots also revealed effects of the introduced mutations on HERD-1 migration through SDS-PAGE gels: phospho-dead HERD-1 migrated significantly slower than HERD-1 from wild-type animals, whereas the phospho-mimic version of HERD-1 migrated between wild-type and phospho-dead and phospho-dead version. We currently cannot explain the magnitude of these migration effects.

In addition, we performed immunoprecipitation analyses (IP) followed by mass spectrometry for HA-tagged HERD-1 and HERD-1 phospho-dead mutant. Results for the wild-type HERD-1 reproduced previously published data (Zhao *et al*., 2024) (Fig. EV4A). In the HERD-1 phospho-dead mutant IP we found that Z granule components LOTR-1, WAGO-4, PID-2, PID-5, PRG-1, ZNFX-1, MIP-2, and P granule component MIP-1 were more enriched than in HERD-1 IP(Huang *et al*., 2025)(Fig. EV4B,C), suggesting that HERD-1 phosphorylation is involved in the regulation of protein-protein interactions within the granules.

We conclude that DEPS-1 and HERD-1 phosphorylation are important for proper ZNFX-1 granule morphology and protein-protein interactions within these granules. In addition, independent of phosphorylation status, mutations in the C-terminal region of both proteins affect PGL-1 granule intensity and the ratio between ZNFX-1 and PGL-1 granules number.

### Loss of DEPS-1 and HERD-1 phosphorylation causes enhanced RNAi inheritance

Next, we asked how changes in DEPS-1 and HERD-1 phosphorylation could affect the inheritance of small RNA-mediated silencing. To address this question, we performed *gfp* RNAi in wild-type and mutant strains carrying a GFP::H4 reporter (Fig. 5A). All strains also expressed PGL-1::mTagRFP, which we used to normalize GFP fluorescence and thus control for positional and size effects of the germline. As reported earlier, PGL-1 signal is higher in the DEPS-1;HERD-1 phospho-dead double mutant than in wild-type, leading to a lower Relative fluorescence intensity value in the mutant. Because we compared control RNAi and *gfp* RNAi within one genotype, this effect did not affect our conclusions. In the DEPS-1;HERD-1 phospho-dead double mutant, the GFP::H4 reporter was normally silenced in the parental (P0) generation, and silencing was transmitted to the next generation (F1) (Fig. 5B). By the third generation, both the wild-type and the mutant began to gain GFP signal. However, while wild-type animals fully restored GFP expression by generation 5, the double-mutant population reached it only one generation later. These results indicate that phosphorylation of DEPS-1 and HERD-1 limits the inheritance of GFP silencing through RNAi. We note that this does not phenocopy *pid-4^-/-^;pid-5^-/-^* mutants, as these were shown to be defective in RNAi inheritance(Placentino *et al*., 2021). Thus, loss of PID-4 and PID-5 appears to affect more than only the phosphorylation of DEPS-1 and HERD-1.

**Figure 5.**
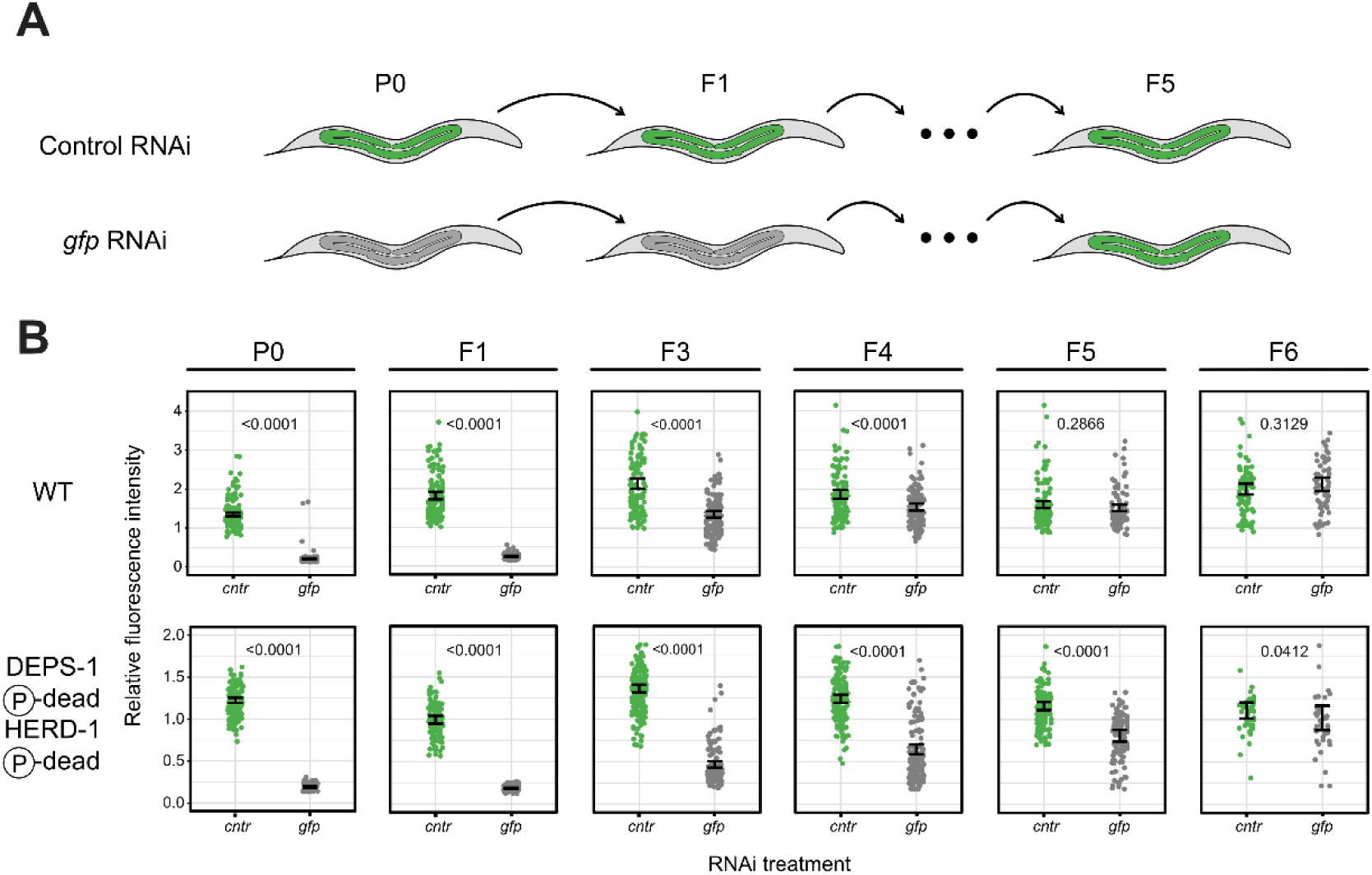
*gfp* silencing inheritance is prolonged in DEPS-1 and HERD-1 phospho-dead double mutant. **(A)** Scheme depicting *gfp* RNAi inheritance in *C. elegans* wild-type. Green represents germline expression of GFP::H4 reporter and gray – silenced reporter. In wild-type animals, silencing is transmitted for 3-5 generations. **(B)** Inheritance of *gfp* RNAi-induced silencing. Relative fluorescence intensity was calculated as GFP signal normalized to RFP signal for each animal. PGL-1 signal is higher in the DEPS-1;HERD-1 phospho-dead double mutant than in wild-type, leading to a lower Relative fluorescence intensity value in the mutant. P0 animals were treated with control RNAi (green) or *gfp* RNAi (gray). F1-F6 progeny were cultured on OP50 plates. P0 and F1, F3, F4, F5, F6 progeny were scored for GFP silencing. Each dot represents an individual animal. Error bars show 95% confidence intervals for the mean. Sample size = ∼60 animals per condition. Statistical significance was assessed by testing contrasts in Gaussian linear models.

### DEPS-1 bridges ZNFX-1 and HERD-1 within Z granules

Although several mutants affecting Z granules, including *pid-2^-/-^*, *lotr-1^-/-^* and *herd-1^-/-^*, have been described(Placentino *et al*., 2021; Wan *et al*., 2021; Marnik *et al*., 2022; Zhao *et al*., 2024), the protein-protein interaction network within Z granules remains poorly understood. We used the most highly enriched proteins from the HERD-1 co-immunoprecipitation experiment (Fig. EV4A) and screened for interactions with HERD-1 using high-throughput AlphaFold2-Multimer predictions (Fig. EV5A)(Hohmann *et al*., 2026). This screen, which uses AlphaFold2 Multimer coupled to MMseqs2, returned no confident predictions. However, predicting the same interactions with AlphaFold3 yielded a high-confidence model for a HERD-1/DEPS-1 heterodimer (Fig. EV5B).

At the same time, we also performed a similar screening with the major Z granule protein ZNFX-1 and used proteins that were identified in a previously published ZNFX-1 TurboID dataset as prey (Zhao *et al*., 2024)(Fig. EV4A). Since ZNFX-1 is a large, multi-domain protein, we divided it based on its AlphaFold model into individual subdomains (Fig. EV5C). Here, the AlphaFold2 pipeline predicted an interaction between DEPS-1 and the N-terminal domain of ZNFX-1 (residues 1-190), and AlphaFold3 models converged on the same interaction interface (Fig. 6A,EV5D). Since this domain has not been annotated previously, we used Foldseek (van Kempen *et al*., 2024) and HHpred (Zimmermann *et al*., 2018), which revealed low sequence similarity but a high structural similarity of this region to MID domains typically found in Argonaute proteins (Fig. EV6A). We therefore refer to this domain as MID-like. In contrast to canonical MID domains, this MID-like domain of ZNFX-1 is highly negatively charged, making it unlikely that this domain retains RNA-binding function (Fig. EV6B). Since both HERD-1 and ZNFX-1 bind to DEPS-1 on different surfaces, we predicted the HERD-1/DEPS-1/ZNFX-1 trimer using AlphaFold3, which resulted in five models that agreed with the previously predicted dimers (Fig. 6B,EV6C).

**Figure 6.**
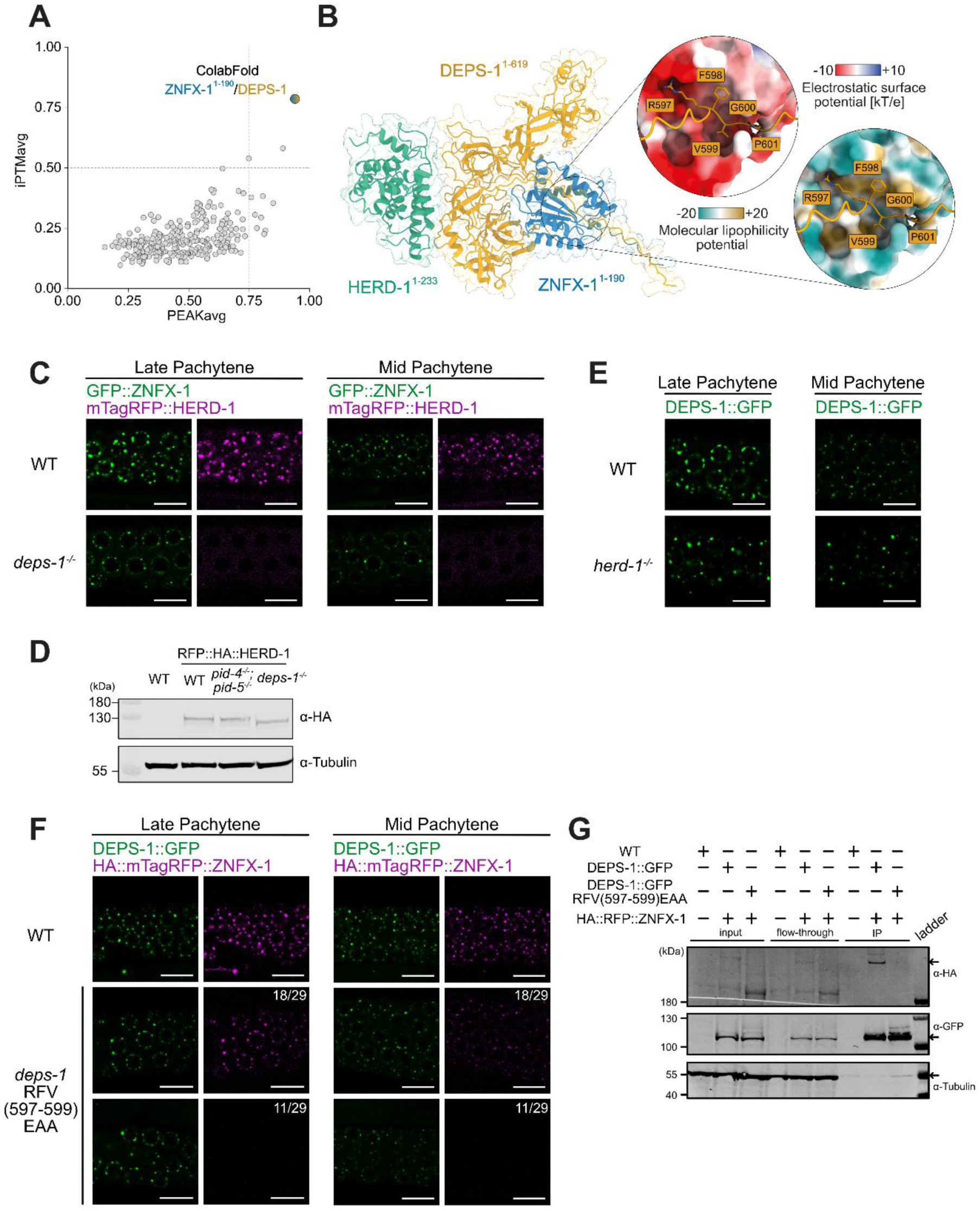
DEPS-1 forms a complex with HERD-1 and ZNFX-1. **(A)** Results of HT-ColabFold screening of ZNFX-1 divided into individual subdomains against proteins previously identified in a ZNFX-1 TurboID (Zhao *et al*., 2024). Dashed lines indicate the thresholds at 0.5 for the iPTM and 0.75 for the PEAKavg. **(B)** AlphaFold3 prediction of full-length DEPS-1 (residues 1-619, yellow) with the N-terminal region of HERD-1 (residues 1-233, green) and ZNFX-1 (residues 1-190, blue). Zoom-in shows the interaction interface between the C-terminal disordered region of DEPS-1 (residues 597-601, yellow) with the highly negatively charged groove on ZNFX-1 (residues 1-190). **(C,E,F)** Representative fluorescence microscopy of GFP::ZNFX-1, PGL-1::mTagRFP, DEPS-1::GFP and HA::mTagRFP::ZNFX-1 in Mid and Late Pachytene regions of the same young adult animals in the indicated backgrounds. n = 8-11 biological replicates. Scale bars, 20 μm. **(D)** Immunoblot analysis of RFP::HA-tagged HERD-1 in indicated mutants. Whole-worm protein extracts from gravid adult hermaphrodites were resolved by SDS-PAGE and probed with anti-HA antibody. α-tubulin served as a loading control. **(G)** Western blot of DEPS-1 and ZNFX-1 after DEPS-1::GFP immunoprecipitation (IP) with GFP-Trap magnetic agarose. Anti-HA antibodies were used to detect HA::mTagRFP::ZNFX-1, anti-GFP – DEPS-1::GFP. α-tubulin served as a loading control; wild-type worms were used as a negative antibody staining control. Arrows indicate proteins of interest level. n = 2.

In all models, the N-terminal region of HERD-1 (residues 1-233) forms an extensive interface with DEPS-1 (residues 315-407). In addition, a short motif in the C-terminal disordered region of DEPS-1 (residues 597-601) binds into this negatively charged groove on the MID-like domain of ZNFX-1 (Fig. 6B). This motif is also in proximity to the DEPS-1 phosphorylation sites T606 and S608. Although the role of phosphorylation in this interaction remains unclear, the addition of negatively charged groups near the site that interacts with the negatively charged MID-like domain might interfere with binding due to charge repulsion. Notably, neither AlphaFold2 Multimer nor AlphaFold3 predicted an interaction between ZNFX-1 and HERD-1, leading us to conclude that DEPS-1 might act as a bridge between the two proteins.

To test this idea, we examined the granular localization of ZNFX-1 and HERD-1 in the absence of DEPS-1. In *deps-1^-/-^* mutants, ZNFX-1 granule intensity was reduced, and HERD-1 completely lost its granular localization, while protein levels remained unchanged (Fig. 6C,D,EV7A). We also observed an increase in DEPS-1 granule intensity, but not ZNFX-1, in the *herd-1^-/-^* mutant (Fig. 6E,7D,EV7B,EV8E).

**Figure 7.**
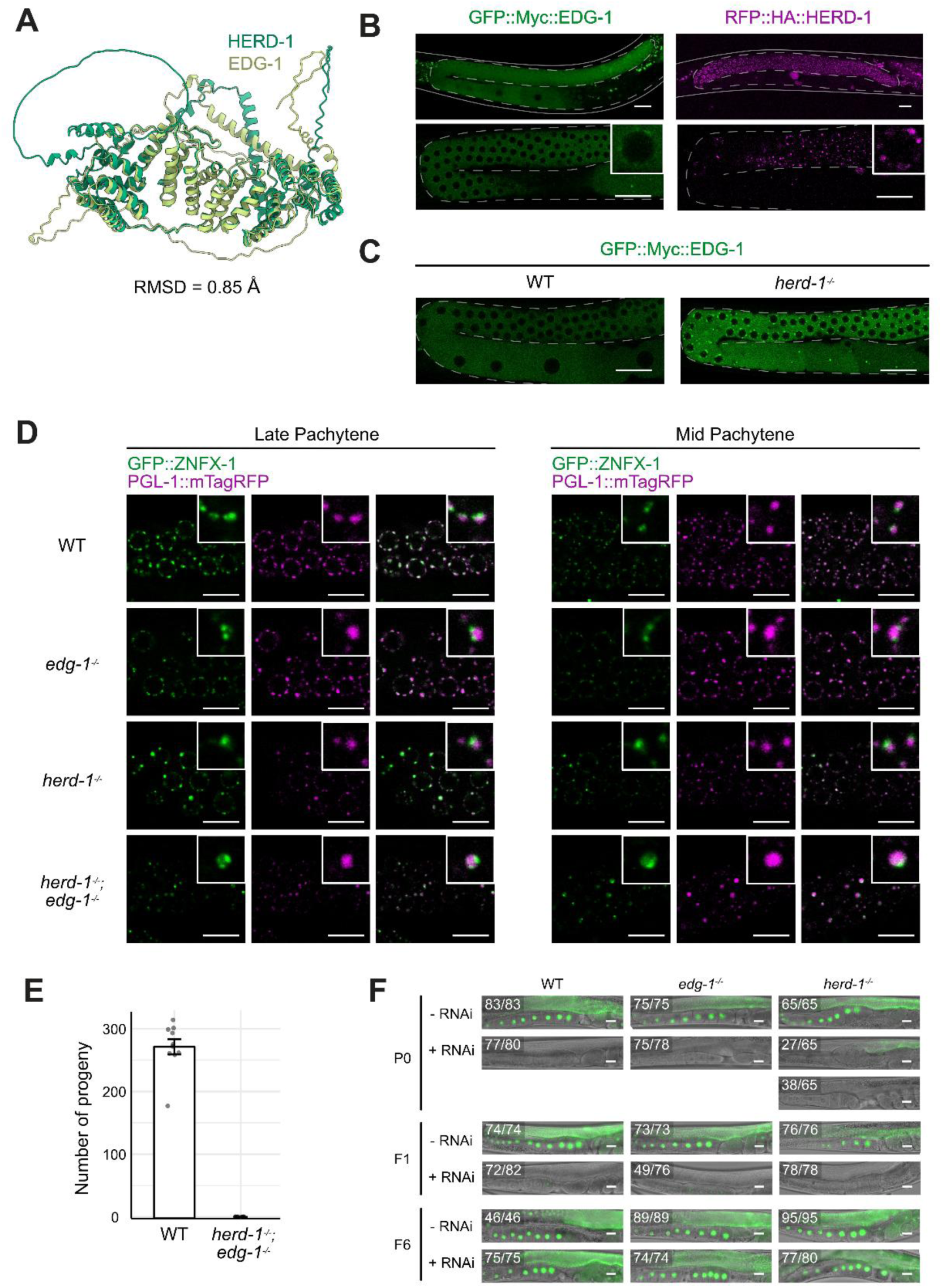
EDG-1 partially compensates for the absence of HERD-1. **(A)** Comparison of HERD-1 (dark green) and EDG-1 (yellow green) structures predicted by AlphaFold3. **(B)** Representative fluorescence microscopy of GFP::Myc::EDG-1 and RFP::HA::HERD-1 in the germline of young adult animals with different zoom-ins. Dashed lines represent the germline boundary. Images are representative of three independent biological replicates. Scale bars, 20 μm. **(C)** Representative fluorescence microscopy of GFP::Myc::EDG-1 in young adult animals in the indicated mutant backgrounds. Dashed lines represent the germline boundary. Images are representative of three independent biological replicates. Scale bars, 20 μm. **(D)** Representative fluorescence microscopy of GFP::ZNFX-1 and PGL-1::mTagRFP in Mid and Late Pachytene regions of the same young adult animals in the indicated backgrounds. Zoom-in of one granule shown in the white box. Scale bars, 10 μm. **(E)** Inheritance of *gfp* RNAi-induced silencing in the indicated mutant backgrounds. Representative fluorescence images of P0, F1 and F6 animals expressing GFP::H4 in the germline. P0 animals were exposed to *gfp* RNAi (+ RNAi) or empty vector control (-RNAi). F1-F6 progeny were cultured on OP50 plates. Numbers indicate animals with the depicted phenotype out of the total scored. Scale bar, 20 μm.

Next, we mutated DEPS-1 residues RFV(597-599) to EAA, to disrupt DEPS-1 interaction with ZNFX-1. In this case, DEPS-1 and ZNFX-1 granular signals were significantly reduced, and in around 30% of the animals we did not detect ZNFX-1 signal at all (Fig. 6F,EV7C,D). In worms that had detectable DEPS-1 and ZNFX-1 signals, proteins showed a tendency to be separate, but did not reach significance (Fig. EV7E,F). Despite its signal reduction in microscopy, the DEPS-1 RFV(597-599)EAA mutation didn’t affect DEPS-1 protein abundance as judged by Western blotting, while ZNFX-1 was highly destabilized (Fig. 6G). We also could not detect ZNFX-1 in DEPS-1 RFV(597-599)EAA immunoprecipitation. HERD-1 signal was reduced in DEPS-1 RFV(597-599)EAA, but the effect was less strong than for ZNFX-1 (Fig. EV7G,H).

Taken together, these findings are consistent with the idea that DEPS-1 plays a central role in Z granule organization by simultaneously binding HERD-1 and ZNFX-1, thereby linking these components within the granule.

### HERD-1 and EDG-1 are partially redundant proteins

We noticed that the HERD-1 and EDG-1 proteins are paralogs that share a high degree of structural and sequence similarity (77% identity, 87% similarity) (Fig. 7A,EV7A,B). Interestingly, EDG-1 had previously been reported as DEPS-1 interactor, and *edg-1* RNAi caused enlargement of DEPS-1 granules (Suen *et al*., 2020), as we observed in *herd-1^-/-^* mutants (Fig. 6E,EV7B). Because of the high sequence homology between the two genes, the RNAi result for *edg-1* could in fact have been caused by HERD-1 loss, or by joint loss of both proteins(Suto *et al*., 2026).

To address this, we started by looking at the localization of EDG-1, N-terminally tagged with GFP-Myc, and HERD-1, N-terminally tagged with RFP-HA. We found that HERD-1 and EDG-1 were both expressed in the germline but had distinct subcellular localizations: HERD-1 was concentrated in granules, whereas EDG-1 was primarily cytoplasmic with only a faint presence in germ granules (Fig. 7B). Noticeably, in absence of HERD-1, EDG-1 acquired pronounced granular localization, suggesting that it may (partially) compensate for the absence of HERD-1 (Fig. 7C). We note that tagging HERD-1 with a fluorescent protein also promoted EDG-1 granular localization, suggesting that the RFP tag perturbed HERD-1 function and/or stability (Fig. EV8C).

To test functional redundancy between HERD-1 and EDG-1, we generated an *edg-1^-/-^* null allele to compare it with the previously described *herd-1^-/-^* allele. In our experiments, the ratio of ZNFX-1 and PGL-1 granule numbers was reduced in *herd-1^-/-^* mutants, but not in *edg-1^-/-^* mutants (Fig. EV8D). In contrast, the intensities of ZNFX-1 and PGL-1 granules in both single mutants were comparable to wild-type (Fig. 7D, EV8E). In addition, ZNFX-1 and PGL-1 signals remained separated in both single mutants. We noted that this is different from what has been reported before by Zhao et al (Zhao *et al*., 2024), who reported ZNFX-1 and PGL-1 granule fusion upon loss of HERD-1. Currently, we cannot explain this difference (see Discussion). The majority of *herd-1^-/-^ ;edg-1^-/-^* double mutant animals failed to develop a germline and were sterile (Fig. 7E). The few *herd-1^-/-^;edg-1^-/-^* double mutant animals that did develop a germline showed partial P and Z granule mixing (Fig. 7D). Together these results indicate that HERD-1/EDG-1 function is essential for germ cell development and that any result on single mutant strains represents a hypomorphic situation. Absence of both HERD-1 and EDG-1 may promote P and Z granule fusion, but the very limited number of animals suitable for analyses prevents a strong conclusion on this aspect.

Finally, we examined the impact of *herd-1^-/-^* and *edg-1^-/-^* mutations on RNAi inheritance using *gfp* RNAi in strains that carry a GFP::H4 reporter(Schreier *et al*., 2025). We did not use the RNAi system we employed in Fig. 5 because, for unknown reasons, the PGL-1 reporter in that system triggered sterility in *herd-1^-/-^* background. Based on the GFP::H4 reporter expression level, we counted the number of animals within two categories: “GFP-expressing” and “silenced”. Consistent with previously published observations (Zhao *et al*., 2024), the P0 generation displayed incomplete silencing of the GFP reporter in *herd-1^-/-^* mutants, whereas the F1 generation was fully silenced (Fig. 7F). This defect in the P0 was not observed in *edg-1^-/-^* mutants. In contrast to the work by Zhao et al (Zhao *et al*., 2024), all strains expressed GFP at the control level by the sixth generation, and thus, no extended RNAi inheritance was observed in *herd-1^-/-^* mutants in our hands (see Discussion).

## DISCUSSION

Over the past years it has become clear that the landscape of germ granules in the nematode germline is complex. Multiple apparently distinct granules have been identified based on interaction and proximity studies and effects of granules on notably RNAi-related processes have been noted. Nevertheless, it remains largely unclear to what extent germ granules are formed to drive specific processes or whether they arise as a consequence of highly abundant transcripts and proteins(Seydoux, 2018). It is thus important to identify potential regulatory mechanisms that can be coupled directly to granule formation and properties, as this will provide a means to recognize cause and consequence of processes related to RNAi and germ granules.

We identify a casein kinase Iα (CKIα) protein, KIN-19, that is present in Z granules and whose presence in Z granules is linked to the phosphorylation of at least two Z granule proteins: HERD-1 and DEPS-1. We also show that blocking of the phosphorylation of specifically these proteins results in a growth of Z granules. This is accompanied by mildly prolonged RNAi inheritance. While we cannot identify the precise molecular processes that lie at the basis of these phenotypes, our results do indicate that post-transcriptional manipulation of granule components affects RNAi, consistent with the possibility that granules contribute functionality to the RNAi response. Another casein kinase family protein, CK2 (Zhang *et al*., 2024) has also been linked to RNAi-like pathways. It was proposed that CK2 acts in the nucleus upstream of piRNA transcription, so not within the context of the here-studied germ granules.

Interestingly, loss of KIN-19 from Z granules affected the granular localization of 14-3-3 protein PAR-5, even though PAR-5 itself is not found within these granules. We found PAR-5 granules to be located close to Z granules, and hypothesize that, being a 14-3-3 protein, PAR-5 may interact with phospho-epitopes on the surface of Z and possibly other granules. As such, it may play a role in positioning specific germ granules relative to each other. This idea will require extensive experimentation to test. As PAR-5 is an essential protein, refined studies aimed specifically at PAR-5 that is present in the germ granules will be required.

We found that reduced amounts of KIN-19 in Z granules lead to the hypo-phosphorylation of Z granule components DEPS-1 and HERD-1. Mutations that prevent DEPS-1 and HERD-1 phosphorylation are accompanied by increased Z granule size, as assessed by ZNFX-1 protein abundance. It is possible that this relates to the enhanced RNAi inheritance that we observed under these conditions. ZNFX-1 has been proposed to promote inheritance of RNAi by re-positioning the 22G generating machinery towards the 3’ end of the targeted transcript. Enlarged ZNFX-1 foci may reflect enhanced repositioning activities on targeted transcripts, thereby affecting the length of the RNAi inheritance response. However, we note that Z granules were also found to be bigger in *pid-4^-/-^;pid-5^-/-^* mutants, but this was accompanied by loss of RNAi, and not by increased inheritance. This difference highlights that the effects of PID-4 and PID-5 extend beyond KIN-19-associated phosphorylation of HERD-1 and DEPS-1 and that the observed phenotypes can presently not yet be directly coupled to effects on individual proteins.

Both HERD-1 and DEPS-1 phosphorylation were found to be KIN-19-dependent, and we also provide evidence that these two proteins may interact with each other. Notably, DEPS-1 has a strong effect on HERD-1. Without DEPS-1, HERD-1 does not localize to Z granules (or to any germ granule), implicating DEPS-1 as a factor that recruits HERD-1 to Z granules. HERD-1 protein levels are not affected by DEPS-1 loss, but interestingly, HERD-1 migration through SDS-PAGE gels is affected: HERD-1 from *deps-1*^-/-^ mutant extracts migrates faster than HERD-1 from wild-type animals. Given that in other experiments we found that phospho-dead HERD-1 migrates slower than wild-type HERD-1, this could indicate that HERD-1 may be hyper-phosphorylated in the absence of DEPS-1, possibly by being exposed to other kinases besides KIN-19 (CKIα). Interestingly, EDG-1, despite high sequence similarity to HERD-1, lacks KIN-19-dependent phosphorylation sites. Whether EDG-1 does become a CKIα substrate in the absence of HERD-1 (a condition in which EDG-1 becomes germ granule-enriched) is currently unclear.

Besides interacting with HERD-1, we provide evidence that DEPS-1 may also bind ZNFX-1. Furthermore, DEPS-1 is known to bind the PIWI protein PRG-1. This starts to draw a scenario in which DEPS-1 may bring PRG-1, with its target, in proximity to ZNFX-1, helping it to stimulate 22G RNA production. In HERD-1 immunoprecipitation, we also found secondary Argonaute such as WAGO-1, WAGO-3 and WAGO-4, suggesting that a similar principle could work for them, as we previously showed that ZNFX-1 acts downstream of WAGO-3-mediated inheritance of RNAi(Schreier *et al*., 2025). PRG-1 and DEPS-1 were also found in the immunoprecipitation of ZNFX-1-interacting protein LOTR-1, indicating that it is also involved in communication between granules (Marnik *et al*., 2022). Interestingly, loss of PRG-1 has been found to enhance RNAi inheritance (Shukla, Perales and Kennedy, 2021; Priyadarshini *et al*., 2022). Possibly, in the absence of PRG-1 the HERD-1/DEPS-1/ZNFX-1 hub that we identify may sustain other RNAi responses better, allowing enhanced inheritance of RNAi. Finally, the fact that the phosphorylation sites that we identify in DEPS-1 are close to the predicted ZNFX-1 interaction site suggests that this proposed bridging function of DEPS-1 may be regulated by phosphorylation.

Possibly, this idea may also be relevant to the previously reported finding of enhanced RNAi inheritance in *herd-1^-/-^* mutants, accompanied by the merging of Z and P granules (Zhao *et al*., 2024). In our experiments, however, we did not observe such effects in *herd-1^-/-^* mutants. The effect is also absent in the recently reported *herd-1* mutant that lacks 685 base pairs of exon four (Suto *et al*., 2026). Currently, we cannot explain this difference. We did observe mild enhancement of RNAi inheritance in the phospho-dead mutants of HERD-1 and DEPS-1, supporting the idea that these proteins indeed affect this process. Possibly, the effects of the *herd-1^-/-^* mutation are subject to strain-specific modifiers of the RNAi response. For instance, we can envisage that expression levels of the HERD-1 paralog EDG-1 may differ between strains, thereby modifying to what extent loss of HERD-1 may be compensated by EDG-1. This idea would be supported by the fact that we did see a tendency to granule merging in the absence of both HERD-1 and EDG-1, but our analysis of this strain was complicated by the fact that the vast majority of double mutant animals failed to develop a germline. This phenotype resembles the sterility that we recently reported in situations where WAGO-3 becomes misloaded with histone mRNA-targeting 22G RNAs(Isolehto *et al*., 2025), raising the possibility that the *herd-1^-/-^;edg-1^-/-^* sterility may be driven by the misloading of Argonaute proteins. Further work will be required to resolve this.

## METHODS

### Worm culture

*C. elegans* strains were maintained on nematode growth medium (NGM) agar plates seeded with *E. coli* OP50 at 20°C under standard laboratory conditions(Brenner, 1974). The Bristol N2 strain served as the wild-type reference throughout this study. A complete list of strains is provided in the Reagents and Tools Table.

### CRISPR-CasG genome editing

For plasmid-encoded Cas9 injections, protospacer sequences were identified using CRISPOR (http://crispor.tefor.net) (*Haeussler et al*., 2016) and inserted into pRK2411 (a derivative of pDD162,#47549, Addgene) through site-directed, ligase-independent mutagenesis (SLIM)(Chiu *et al*., 2004). SLIM products were introduced into DH5α competent cells (Invitrogen™) and grown on LB agar containing 100 μg/ml ampicillin. We injected 50 ng/µl Cas9 + co-conversion guide RNA plasmid, 50 ng/µl gene of interest guide RNA plasmid, 750 nM co-conversion single-stranded oligodeoxynucleotide (ssODN) donor (IDT) and 750 nM gene of interest ssODN donor (IDT)(Schreier *et al*., 2022).

For Cas9 protein injections, crRNAs were designed using the Integrated DNA Technologies CRISPR–Cas9 platform (https://eu.idtdna.com/site/order/designtool/index/CRISPR_CUSTOM). Strains were created by microinjecting 2.5µg/µl recombinant Cas9 protein (produced in-house), 500nM of ssDNA (IDT) or PCR product as repair template, 440 nM co-conversion ssODN donor (IDT), 42.5µM tracrRNA (IDT), 12µM dpy-10 guide RNA (IDT) and 30µM targeted gene guide RNA (IDT) as described previously(Paix *et al*., 2017). For HERD-1 and EDG-1 tagging with fluorescent proteins, HiFi Cas9 Nuclease V3 (1081060, IDT) and ssODN templates were used. To obtain ssODN templates, one of the primers in the PCR mixture had a 5′-phosphate modification, and the PCR product was treated with λ-exonuclease (M0262, NEB)(Eroglu, Yu and Derry, 2023).

Genome editing used *dpy-10(cnc4)* co-conversion approaches(Arribere *et al*., 2014). Guide RNA sequences and repair templates are provided in the Reagents and Tools Table. F1 offspring were identified by PCR and validated through Sanger sequencing. Mutants were outcrossed twice with wild-type N2 animals to eliminate potential off-target mutations.

### Microscopy

For live imaging, 20–30 animals were washed in 100 µl M9 buffer and then transferred to 50 µl M9 buffer containing 40 mM NaN_3_ (S002, Sigma-Aldrich) on a coverslip. After worms stopped moving, a glass slide with a freshly prepared 2% (w/v) agarose pad in water was placed on top of the coverslip. Samples were imaged on a TCS SP5 confocal microscope with a 40x/1.3 oil objective. Images used to identify the distance between granules were obtained with a Zeiss LSM900 microscope with a 63x/1.4 oil-immersion objective.

For RNAi inheritance, one sample slide contained at least 75 adult hermaphrodites. Animals were imaged using a THUNDER Imager (Leica) inverted widefield microscope equipped with a 10x/0.32 dry objective for tile scan acquisition or 40x/0.95 dry objective for individual animal acquisition.

### Immunostaining

Adult animals were dissected in Egg buffer (25 mM HEPES, pH 7.4, 118 mM NaCl, 48 mM KCl, 2 mM EDTA, 0.5 mM EGTA, 10mM NaN_3_) supplemented with 0.1% Tween-20 and fixed in 0.3% formaldehyde diluted in Egg buffer. Following freeze-cracking on dry ice, specimens were transferred to pre-chilled acetone and incubated for 10 min. Germ lines were incubated with anti-GFP nanobody abberior STAR GREEN 1:500 dilution (N0304-AbGREEN-L, NanoTag Biotechnologies), primary anti-Myc antibody at 1:1000 dilution (2276S, Cell Signaling Technology), and secondary anti-mouse IgG Alexa Fluor 555 at 1:1000 dilution (A11029, Thermo Fisher). Samples were mounted using ibidi mounting medium containing DAPI (50011, ibidi).

### Image processing

For image processing, ImageJ v.1.54t was used(Schindelin *et al*., 2012). Images were prepared for publishing in Omero(Besson *et al*., 2019). Data plotting was performed in R (version 4.3.2) using the ggplot2 package. Detailed custom FIJI Macroses for image analyses are attached in OMERO to the corresponding folder.

To assess the degree of colocalization between two granules, AiryScan-processed images were used. First, we cropped an area containing 1 nucleus within the pachytene zone of the gonad arm. The ImageJ “Find Maxima…” function was used to identify the pixel of peak intensity within each granule. The distance between the peak-intensity pixels of the closest granules labeled for different proteins was then measured. Distance in pixels was converted to microns. Germ granules surrounding 3–4 nuclei per animal were quantified, using 3–5 animals per experiment. The mean inter-granule distance was calculated for each worm, and statistical significance was assessed with Welch’s tests or Dunnett’s test when correction for multiple experiments was needed.

To determine granule number, we cropped an area containing the whole pachytene zone of the gonad arm and applied the “Find Maxima…” function. For PAR-5 analyses, the blurred image was subtracted from the original to eliminate cytoplasmic signal and analyze only granular signal. The same prominence value was used for a protein of interest across all images. 10–17 animals were used per experiment. For plotting, the ratio of granules of interest to PGL-1 granules was used. P values were calculated using Welch’s test or Dunnett’s test when correction for multiple experiments was needed.

To determine maximum intensity of germ granules, we cropped an area containing the whole pachytene zone of the gonad arm. For ZNFX-1 channel overall intensity, a local-mean filter (radius = 2 px) and a modest Gaussian blur (σ = 2) were applied, and a fixed intensity threshold (40–255) was used to generate a binary mask. After median filtering (radius = 2) and watershed segmentation, particles larger than 0.02 µm² were identified and stored as regions of interest ROIs. ROIs were applied to the original KIN-19 and ZNFX-1 channels to measure max intensity. The same threshold values were used for a protein of interest across all images. 6-7 animals were used per experiment. For each worm we calculated the mean of the log₂-transformed maximum intensity of all granules. P values were calculated using a linear mixed model.

For granule intensity analysis, the early and late pachytene regions of the germline were analyzed separately, as Z granule protein intensity differs between these regions. For each channel, a local-mean filter (radius = 2 pixels) and a modest Gaussian blur (σ = 2) were applied to reduce high-frequency noise. A fixed intensity threshold was used to create a binary mask, which was subsequently median-filtered (radius = 2) and subjected to watershed segmentation to separate touching objects. Particles larger than 0.02 µm² were identified and stored as ROIs. These ROIs were overlaid on the unblurred duplicate of the corresponding channel, and the raw integrated intensity was measured. The same threshold values were used for a protein of interest across all images. 10–17 animals were used per experiment. P values were calculated using a linear mixed model, followed by Holm’s correction for multiple comparisons.

For RNAi inheritance in strains without PGL-1::mTagRFP expression, the number of animals within two categories was quantified: “GFP-expressing” (fluorescence intensity comparable to non-RNAi controls) or “silenced” (complete absence of detectable GFP or barely detectable fluorescence).

RNAi inheritance in strains expressing PGL-1::mTagRFP was quantified using a custom ImageJ macro, following the previously described approach (Schreier *et al*., 2025). Before analysis, a 100-pixel border was removed from each edge of every image to eliminate artifacts introduced during tile stitching. Individual worm germlines were then segmented based on the RFP fluorescence signal. For each image, the region outside all detected bright objects was defined as a background ROI, and the mean intensity within this region was subtracted from the entire image separately for the GFP and RFP channels, correcting for uneven illumination and background autofluorescence across the stitched tiles. Following automated segmentation, all ROIs were manually inspected, and any objects incorrectly identified as worms were excluded. Integrated and mean fluorescence intensities were then measured for each validated worm ROI in both channels. To correct for positional effects along the germline, the ratio of integrated GFP to RFP intensity was calculated. This ratio was log2-transformed and analyzed using a Gaussian linear model. Normality of the values was confirmed using Q–Q plots and Shapiro–Wilk tests. Models were fit in R (version 4.3.2) using the *lm* function, and contrast *p*-values with corresponding confidence intervals were obtained using the *emmeans* package (version 2.0.4).

### Immunoprecipitation

Synchronized animals were maintained at 20°C until reaching the young adult stage. Worms were washed with M9 buffer, resuspended in sterile water as 200 μl aliquots, and flash-frozen on dry ice. Frozen worm samples were thawed on ice and resuspended in 500 μL of cold lysis buffer (25 mM Tris-HCl pH 7.5, 150 mM NaCl, 1.5 mM MgCl₂, 1 mM DTT, 0.1% Triton X-100, and cOmplete Mini EDTA-free protease inhibitor cocktail (11836170001, Roche)). Lysates were sonicated with Bioruptor Plus (Hologic Diagenode) for 10 cycles of 30 seconds on and 30 seconds off at high efficiency, with sonication repeated if cells were not completely lysed. Samples were centrifuged at maximum speed for 10 minutes at 4°C, and supernatants were collected. Protein concentrations were determined by BCA assay, and equal amounts of protein were used for each sample. Input samples (20 μL) were removed and stored at −20°C.

For HERD-1::HA and HERD-1::HA(S712A;S713A) immunoprecipitation, 450 μL of lysate was added to the 30 μL of anti-HA magnetic beads (88837, Thermo Fisher). For DEPS-1::GFP and DEPS-1::GFP(RFV(597-599)EAA) immunoprecipitation, lysate was added to the 20 μL of GFP-Trap magnetic agarose (gtma, Proteintech). The mixture was incubated overnight at 4°C with rotation and 20 μL of flow-through was collected and stored at −20°C. Beads were then washed three times with 500 μL of wash buffer (25 mM Tris-HCl pH 7.5, 150 mM NaCl, 1.5 mM MgCl₂, 1 mM DTT, and cOmplete Mini EDTA-free protease inhibitor cocktail (11836170001, Roche)). Proteins were eluted by adding 50 μL of resuspension buffer (25 mM Tris-HCl pH 8.0, 1% SDS) and boiling at 70°C for 10 minutes. Eluates were collected and stored at −20°C until analysis.

### TMT-based phosphoproteomics

For the wild type and the *pid-4^-/-^;pid-5^-/-^* double mutant phosphoproteomics and proteomics synchronized animals were maintained at 20°C until reaching the young adult stage. Worms were washed with M9 buffer, resuspended in sterile water as 200 μl aliquots, and flash-frozen on dry ice.

*C. elegans* were lysed in 6 M urea / 2 M thiourea in 50 mM TRIS pH 8 supplemented with protease and phosphatase inhibitors. Lysates were incubated for 15 min, then 5M NaCl was added (1:10 V:V). Lysates were sonicated using the Bioruptor Plus (10×30 sec, high intensity, with 30 s rest cycles) to enable chromatin extraction. Lysates were centrifuged for 15 min at 16 000 ×g and supernatant concentrations measured using the Bradford assay. Samples were reduced with 2 mM DTT for 45 min and alkylated using 11 mM chloroacetamide for 30 min. Samples were diluted in 4x (V/V) excess of 50 mM Tris buffer, pH 8. Proteins were digested using 1:50 (w:w) MS-grade trypsin (Serva) overnight at RT. Samples were incubated in TFA (0.5% V/V) for 1 h at 4°C and centrifuged for 10 min at 4,000 ×g. Peptide supernatants were purified using C18 Sep–Pak columns (Waters), eluted in 50% acetonitrile and dried using SpeedVac. Peptides (0.2 mg) resuspended in 150 mM HEPES (pH 8.5) / 30% ACN were barcoded with 0.2 mg of acetonitrile-dissolved TMT10plex labels (Thermo Scientific) for 1 h and quenched with 5% hydroxylamine for 15 min (Li *et al*., 2020). An equal aliquot (5% vol) of each sample was mixed and the ratio check was performed as described(Zhang and Elias, 2017). After the ratio check, masses were adjusted treatment-wise, with the assumption of equal mass distributions across all samples. An equal mass of each sample was multiplexed. Samples were diluted with 0.1% TFA to reach < 3% ACN and purified on SepPak columns, eluted with 50% ACN and acidified with 6% TFA. 1/30 (w:w) of the sample was taken for full proteome analysis. Phosphopeptides were enriched using titanium dioxide resin as described (Borisova, Wagner and Beli, 2017). Samples were fractionated using high-pH reversed phase chromatography (Thermo Fisher) using manufacturer’s instructions.

### Liquid chromatography tandem mass spectrometry

For the wild type and the *pid-4^-/-^;pid-5^-/-^* double mutant proteomics, samples were analysed on a quadrupole Orbitrap mass spectrometer (Exploris 480, Thermo Scientific) equipped with a UHPLC system (EASY-nLC 1200, Thermo Scientific). They were loaded onto a C18 reversed-phase column (55 cm length, 75 mm inner diameter, packed in-house with ReproSil-Pur 120 C18-AQ 1.9-mm beads, Dr. Maisch) and eluted with a gradient from 2.4 to 33.6% acetonitrile containing 0.1% formic acid in 120 min. The mass spectrometer was operated in a data-dependent mode, automatically switching between MS and MS2 acquisition. Survey full scan MS spectra (*m*/*z* 300– 1,650, resolution: 60,000, target value: 3 × 10^6^, maximum injection time: 40 ms) were acquired in the Orbitrap. The 20 (proteome) or 15 (phosphoproteome) most intense precursor ions were sequentially isolated, fragmented by higher energy C-trap dissociation (HCD) and scanned in the Orbitrap mass analyser (normalized collision energy: 33%, resolution: 15,000, target value: 1 × 10^5^, maximum injection time: 40 ms, isolation window: 0.8 *m*/*z*). Precursor ions with unassigned charge states, as well as with charge states of +1 or higher than +6, were excluded from fragmentation. Precursor ions already selected for fragmentation were dynamically excluded for 30 s.

For HERD-1::HA and HERD-1::HA(S712A;S713A) immunoprecipitation samples were processed using the SP3 approach (Hughes *et al*., 2019). Proteins were reduced in 5 mM DTT, alkylated in 15 mM iodoacetamide in the dark and quenched in 5 mM DTT. Enzymatic protein digestion was performed using trypsin at 37°C overnight. Following acidification by formic acid, the peptides were purified by solid phase extraction in a C18 StageTip format (Rappsilber, Ishihama and Mann, 2002). Peptides were separated via an in-house packed 30-cm analytical column (inner diameter: 75 μm; ReproSil-Pur 120 C18-AQ 1.9-μm silica particles, Dr. Maisch GmbH) on an EASY-nLC 1000 UHPLC system (Thermo Scientific). The online reverse phase chromatography separation was conducted through a 105-min gradient of 1.6-32% acetonitrile with 0.1% formic acid at a nanoflow rate of 225 nl/min. The eluted peptides were sprayed directly by electrospray ionization into a Q Exactive Plus Orbitrap mass spectrometer (Thermo Scientific). Mass spectrometry was conducted in data-dependent acquisition mode using a top10 method with one full scan (scan range: 300 to 1,650 m/z; resolution: 70,000, target value: 3 × 106, maximum injection time: 20 ms) followed by 10 fragment scans via higher energy collision dissociation (HCD; normalized collision energy: 25%, resolution: 17,500, target value: 1 × 105, maximum injection time: 120 ms, isolation window: 1.8 m/z). Precursor ions of unassigned or +1 charge state were rejected. Additionally, precursor ions already isolated for fragmentation were dynamically excluded for 20 s.

### Mass spectrometry data processing and statistical analysis

Raw data files were analyzed using MaxQuant (Tyanova, Temu and Cox, 2016). For the wild type and the *pid-4^-/-^;pid-5^-/-^* double mutant phosphoproteomics and proteomics analyses site localization probabilities were determined by MaxQuant using the post–translational modification scoring algorithm. Parent ion and MS2 spectra were searched against a reference proteome database containing human protein sequences obtained from UniProtKB using Andromeda search engine (PMID: 21254760). Spectra were searched with a mass tolerance of 6 p.p.m. in MS mode, 20 p.p.m. in HCD MS2 mode, strict trypsin specificity, and allowing up to two miscleavages. Cysteine carbamidomethylation was searched as a fixed modification, whereas protein N– terminal acetylation, methionine oxidation, and phosphorylation (STY; only for phosphoproteomics) were searched as variable modifications. The dataset was filtered based on posterior error probability (PEP) to arrive at a false discovery rate of below 1% estimated using a target–decoy approach (Elias and Gygi, 2007).

For HERD-1::HA and HERD-1::HA(S712A;S713A) immunoprecipitation analyses spectral data were searched against a target-decoy database consisting of the forward and reverse sequences of the HERD-1::HA and the mutant HERD-1::HA (S712A, S713A) bait proteins, the UniProt *C. elegans* (release 2023_02; 28,540 entries) and *E. coli* (release 2023_01; 5,064 entries) reference proteomes and a list of common contaminants. Trypsin/P specificity was assigned. A maximum of 2 missed cleavages were tolerated. Carbamidomethylation of cysteine was set as fixed modification. Methionine oxidation and protein N-terminal acetylation were chosen as variable modifications. The “second peptides” option was switched on. The “match between runs” function was activated. The minimum peptide length was set to 7 amino acids. False discovery rate (FDR) was set to 1% at both peptide and protein levels.

### Statistical analysis of MS data

For the wild type and the *pid-4^-/-^;pid-5^-/-^* double mutant phosphoproteomics and proteomics experiments statistical analysis and MS data visualization were performed using the RStudio software environment (version 2024.04.2). For protein-level analyses, potential contaminants, reverse hits, hits only identified by site and hits with no unique peptides were excluded from the analysis. Phospho-sites were filtered based on localization probability (>75%). Sample intensities were normalized using quantile normalization approach with assumption of equal intensity distribution across all samples (limma package) (Ritchie *et al*., 2015). Phospho-site intensities were normalized by dividing with proteome intensities to account for treatment-induced changes in protein levels. Statistical significance was calculated using a moderated t-test (limma package) (Ritchie *et al*., 2015).

For HERD-1::HA and HERD-1::HA(S712A;S713A) immunoprecipitation experiments the MaxLFQ algorithm (Cox *et al*., 2014) was employed for label-free protein quantification without using its default normalization option. Minimum LFQ ratio count was set to one. Both the unique and razor peptides were used for quantification. Detected *E. coli* proteins, reverse hits, potential contaminants and “only identified by site” protein groups were filtered out. Data normalization was performed on the log-transformed data. Under the assumption that the majority of the detected proteins were non-specific background binders, the distribution of protein LFQ intensities was assessed using kernel density estimation to find the peak density (Rai *et al*., 2018)]. The LFQ intensities of each pull-down were then normalized by adjusting to the point at which the peak density was found. Proteins were further filtered to retain only those detected in at least 2 out of the 4 replicates in either group of each comparison. Following imputation of the missing LFQ intensity values, the statistical significance of the difference between the two groups was assessed using a modified t-statistic (t(SAM, statistical analysis of microarrays) (Tusher, Tibshirani and Chu, 2001) and visualized on a volcano plot. The combined significance threshold (hyperbolic curve) was defined as t0 = 1 and s0 = 1.5.

### Western Blot

For HERD-1 Western blot analysis, 100 adult worms were manually collected and incubated in 1× NuPAGE™ LDS sample buffer (NP0007, Invitrogen) supplemented with 100 mM DTT at 95°C for 30 min. Protein lysates were resolved on NuPAGE 4–12% Bis-Tris gels (MP0321BOX, Thermo Fisher Scientific) using MOPS NuPAGE running buffer (NP0001, Thermo Fisher Scientific), at 120 V, with Prestained Protein Standard (26616, Thermo Fisher Scientific) included for molecular weight reference.

For DEPS-1 Western blot analysis, protein lysates were resolved on NuPAGE 3-8% Tris-Acetate gels (EA0375BOX, Thermo Fisher Scientific) using Tris-Acetate NuPAGE running buffer (LA0041, Thermo Fisher Scientific), at 120 V, with Prestained Protein Standard (26616, Thermo Fisher Scientific) included for molecular weight reference.

Separated proteins were electrotransferred onto Amersham™ Protran® Western blotting nitrocellulose membranes (GE10600002, SigmaAldrich), at 140 V for 1 hour using a Transfer buffer (prepared in-house) containing 15% methanol. Membranes were blocked for 1 hour in PBS supplemented with 5% skim milk and 0.1% Tween-20, then cut into sections based on molecular weight markers. Individual membrane sections were incubated overnight at 4°C with primary antibodies diluted in blocking buffer: mouse anti-HA (1:2000; H3663, Sigma-Aldrich), mouse anti-GFP (1:1000; sc-9996, Santa Cruz) and mouse anti-α-tubulin (1:4000; ab7291, Abcam). Following five washes of 5 min each with PBS containing 0.1% Tween-20 (PBS-T), membranes were incubated for 1 hour with IRDye 800CW-conjugated goat anti-mouse secondary antibody (1:10,000; 926-32211, LI-COR), then washed three additional times with PBS-T. Immunoreactive bands were visualized using a LI-COR Odyssey M imaging system.

### RNAi inheritance

After alkaline hypochlorite treatment of gravid adults, isolated eggs were placed on RNAi plates seeded with HT115(DE3) bacteria expressing empty vector L4440 (Addgene plasmid #1654; gift from Andrew Fire) or RNAi constructs targeting *xfSi255*[*his-c7p::gfp::his-c7::tbb-2 3’UTR + Cbr-unc-11S(+)*](Schreier *et al*., 2025). In the next generations, eggs were transferred to NGM plates seeded with OP50 *E. coli* (no RNAi). After 4 days at 20°C, adults were scored for GFP expression.

### Structure predictions and analysis

Protein interaction screenings with HT-ColabFold were performed as previously described(Hohmann *et al*., 2026). For the evaluation, predictions which had either an iPTMavg above 0.5 or a PEAKavg above 0.75 were manually inspected. Individual protein and protein complex predictions were carried out with AlphaFold3 using AlphaFold Server with the standard settings (Abramson *et al*., 2024). Graphical representations of protein structures were prepared using ChimeraX 1.10.1(Pettersen *et al*., 2021).

### Protein sequence analyses

Protein sequences were aligned using Clustal Omega(Sievers *et al*., 2011). The final figure was generated using ESPript 3.2(Robert and Gouet, 2014).

## Data availability

The mass spectrometry-based proteomics data for the wild type and the *pid-4^-/-^;pid-5^-/-^* double mutant have been deposited to the ProteomeXchange Consortium via the PRIDE (PMID: 39494541) partner repository with the dataset identifier ProteomeXchange: PXD083363.

Log in to the PRIDE website using the following details:

Project accession: PXD083363 Token: dO5WySkcwQxA

The mass spectrometry data for HERD-1::HA and HERD-1::HA(S712A;S713A) have been uploaded to the UCSD Mass Spectrometry Interactive Virtual Environment (MassIVE) server with the dataset identifier ProteomeXchange: PXD083603.

https://proteomecentral.proteomexchange.org/cgi/GetDataset?ID=PXD083603

MassIVE dataset accession ID: MSV000103091

Credentials for access by reviewers are:

ftp://MSV000103091@massive-ftp.ucsd.edu

Username: MSV000103091_reviewer

Password: G!!H3*%Vm79Pe$Nu

Raw microscopy data can be found on a public Omero site: https://omero.imb.uni-mainz.de/webclient/

## Acknowledgements

We thank the members of the Ketting lab for helpful discussions and feedback on the manuscript. Carolyn M. Phillips for generously providing strains. We thank the IMB Media Laboratory and Microscopy Core Facility for consumables and equipment (TCS STED CW SUPER-RESOLUTION, DFG grant 204504610; AF7000, DFG grant 212049334). We thank the microscope core facility at the Faculty of Biology of the JGU for the use of the LSM900, funded by the Ministry of Science and Health of Rhineland-Palatinate and the European Regional Development Fund (ERDF/REACT-EU, Grant No. 84012490). We thank Jiaxuan (Jimmy) Chen and Amitkumar Fulzele for assistance with mass spectrometry analysis. The authors thank Fridolin Kielisch for assistance with Biostatistics analyses. The work was funded by the Deutsche Forschungsgemeinschaft (DFG, German Research Foundation): project numbers 464588647 (CRC1551, Polymer Concepts in Cellular Function), 407023052 (RTG2526-GenEvo), 252386272 (R.F.K.)(Ketting), BE 5342/2-1 – FOR 2800 (Beli). HT-Colabfold screenings were performed using the Life Science Compute Cluster (LiSC) of the University of Vienna.

## Notes

### Competing Interest Statement

The authors have declared no competing interest.

### Summary of Updates

We included a supplemental data file.

