## Supplemental data for "Phosphorylation of Z granule components regulates germ granule organization and RNAi inheritance"

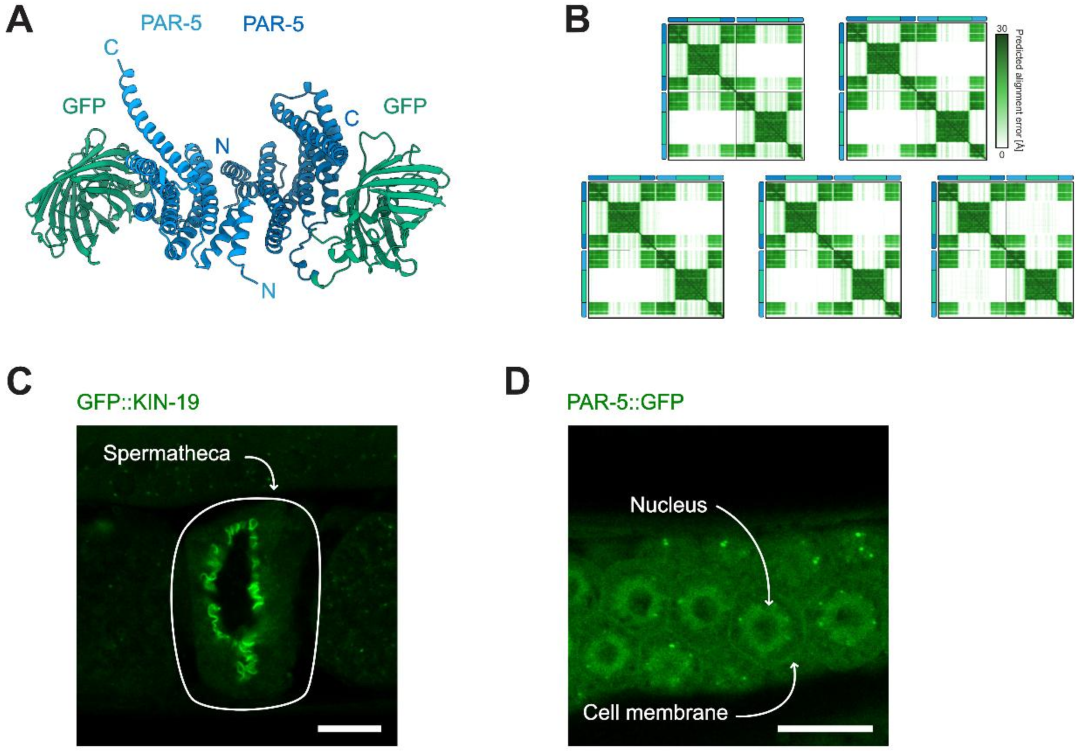


Pshanichnaya, L. *et al.*; Figure EV1

**Figure EV1. Structure of GFP-tagged PAR-5 dimer.**

(A) AlphaFold3 prediction for GFP-tagged PAR-5 dimer. PAR-5 from different monomers are represented in light and dark blue, and GFP is in green.

(B) Predicted Aligned Error (PAE) plots of AlphaFold3 structure from (A). Dashed lines separate protein domains. Color scheme as (A).

(C) Representative fluorescence microscopy of GFP::KIN-19 in the spermatheca. Arrow indicates the spermatheca. Images are representative of three independent biological replicates. Scale bars, 10 µm.

(D) Representative fluorescence microscopy of PAR-5::GFP in the germline. Arrows indicate the nucleus and the cell membrane. Images are representative of three independent biological replicates. Scale bars, 10 µm.

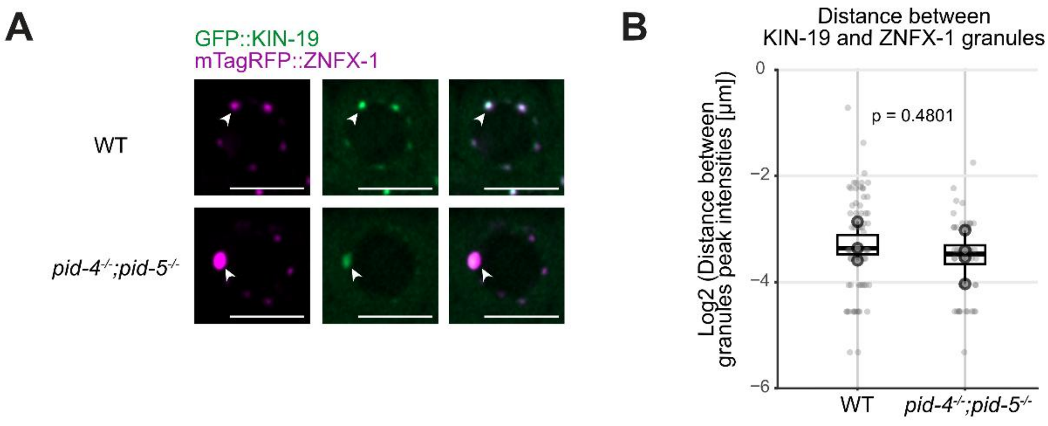


Pshanichnaya, L. *et al.*; Figure EV2

**Figure EV2. KIN-19 retained association with ZNFX-1 in the *pid-4$^{-/-}$;pid-5$^{-/-}$* mutant.**

(A) Representative fluorescence microscopy of one nucleus in the pachytene germline region of young adult hermaphrodites expressing GFP::KIN-19 together with mTagRFP::ZNFX-1 in the indicated mutant backgrounds. Arrows indicate individual perinuclear germ granules. Images are representative of three independent biological replicates. Scale bars, 5 µm.

(B) Quantification of distances between peak intensities of granules shown in (A). Each light dot represents the distance between individual granules; dark dots represent the mean distance between granules within one worm; the box represents the 25th percentile and 75th percentile of the per-worm means, and horizontal bars indicate median values of the per-worm means, whiskers reach the most extreme non-outlier values. For logarithmic data representation, 0 values were replaced with half the pixel size value. Statistical significance was assessed using Welch's t-test. n = 3-4 biological replicates.

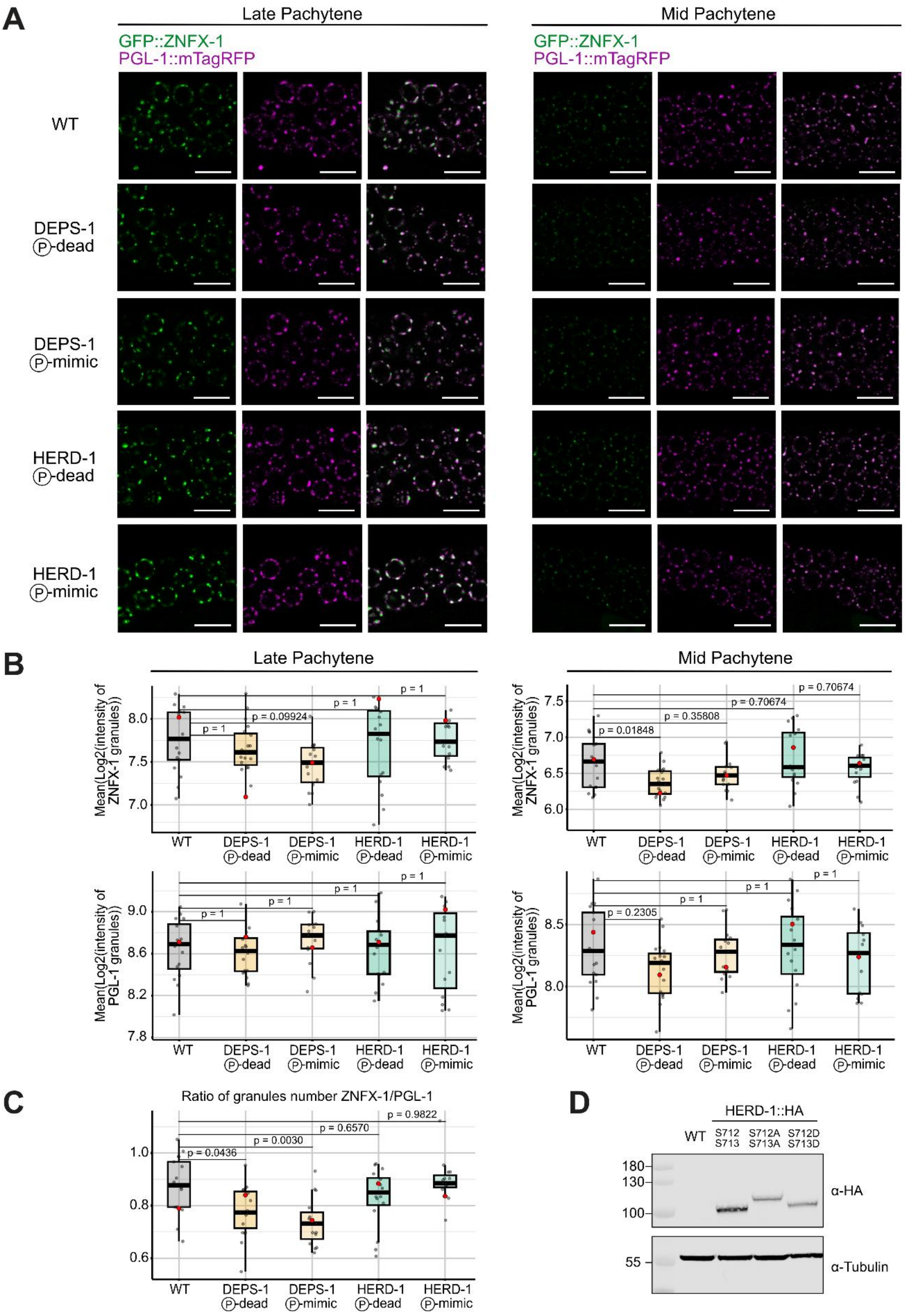


Pshanichnaya, L. *et al.*; Figure EV3

**Figure EV3. Intensity of ZNFX-1 and PGL-1 remains unchanged in DEPS-1 and HERD-1 phosphorylation single mutants.**

(A) Representative fluorescence microscopy of GFP::ZNFX-1 and PGL-1::mTagRFP in Mid and Late Pachytene regions of young adult animals in the indicated backgrounds. DEPS-1 (P)-dead corresponds to DEPS-1 (T606A;S608A) and HERD-1 (P)-dead to HERD-1 (S712A;S713A). DEPS-1 (P)-mimic is DEPS-1 (T606D;S608D) and HERD-1 (P)-mimic is HERD-1 (S712D;S713D). Scale bars, 10 μm.

B) Quantification of ZNFX-1 and PGL-1 granule intensity shown in (A), analyzed separately for the mid and late pachytene regions. Each dot represents the mean log2-transformed granule intensity of an individual animal; boxes indicate the 25th–75th percentile range, and horizontal lines indicate the median, whiskers reach the most extreme non-outlier values. Red dots refer to representative micrographs shown in (B). Statistical significance was assessed using a linear mixed model, with Holm's correction for multiple comparisons. n = 15–19 animals per condition.

(C) Quantification of granule number in the distal region of the germline. The number of ZNFX-1 granules was normalized to the number of PGL-1 granules. Each dot represents the granule ratio for an individual animal; boxes indicate the 25th–75th percentile range, and horizontal lines indicate the median, whiskers reach the most extreme non-outlier values. Statistical significance was assessed using Dunnett's multiple comparison test. n = 15–19 biological replicates.

(D) Immunoblot analysis HA-tagged HERD-1 in indicated mutants. Whole-worm protein extracts from gravid adult hermaphrodites were resolved by SDS-PAGE and probed with anti-HA antibody. α-tubulin served as a loading control.

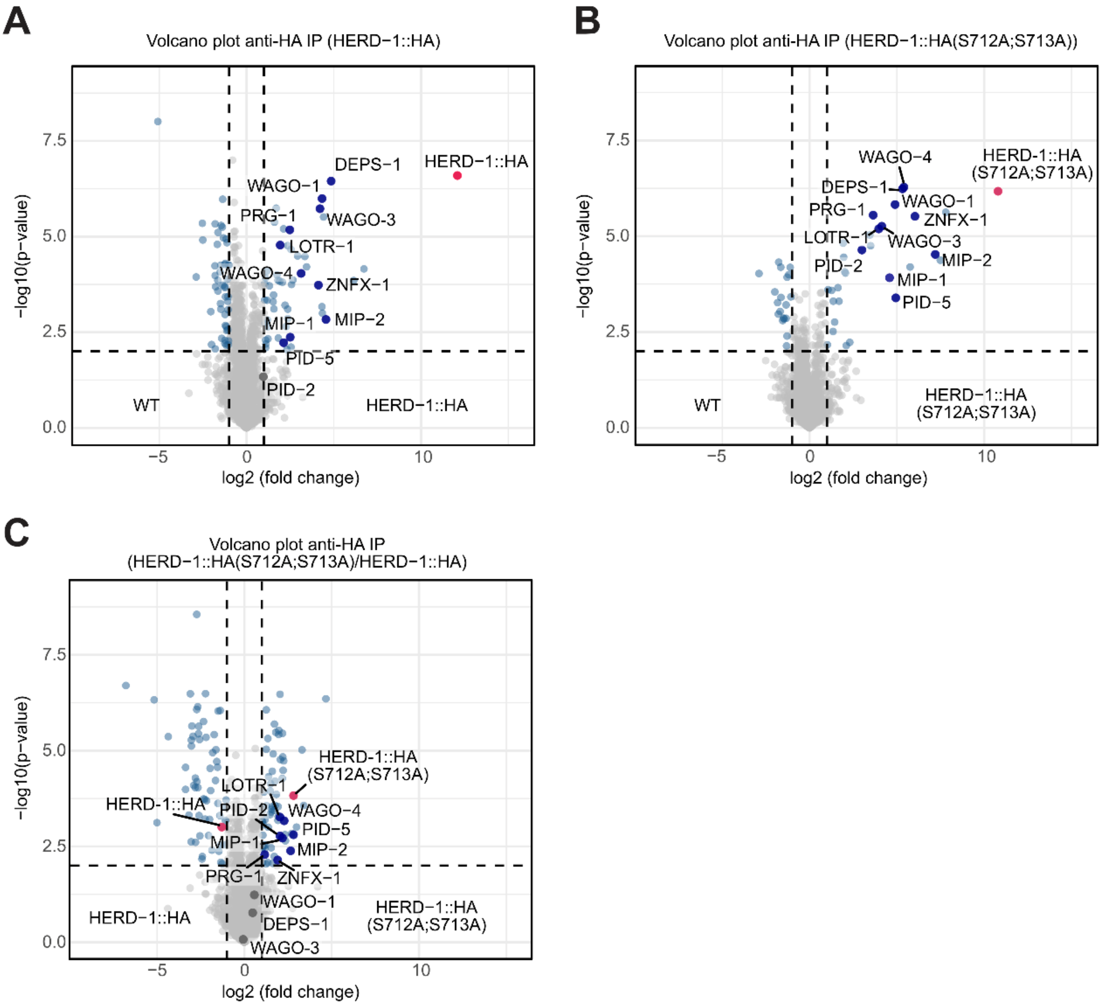


Pshanichnaya, L. *et al.*; Figure EV4

**Figure EV4. HERD-1 pulls down components of Z and P granules.**

(A-C) Volcano plot showing differential protein abundance in HERD-1::HA (A,C) and HERD-1::HA (S712A;S713A) (B,C) immunoprecipitation, quantified using label-free proteomics. Experiments were performed on native protein extracts from young adult hermaphrodites (n = 4 biological replicates). Dashed lines indicate thresholds for significance: p-value = 0.01 and a two-fold change in enrichment. Dots indicate proteins that are enriched or depleted (blue), or detected (grey), bait protein is indicated as a red dot.

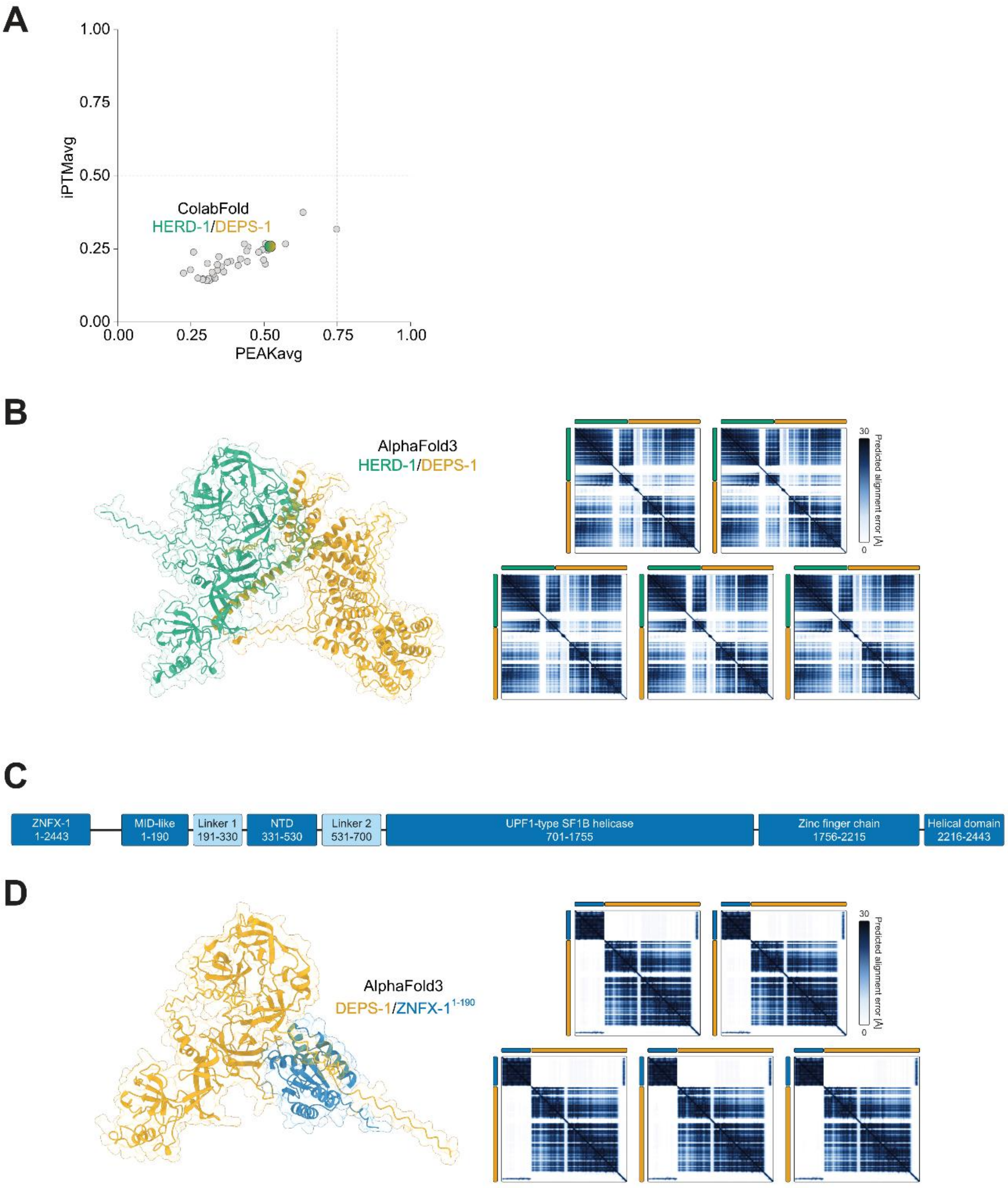


Pshanichnaya, L. *et al.*; Figure EV5

**Figure EV5. AlphaFold3 predictions of interactions within Z granule.**

(A) Results of HT-ColabFold screening of HERD-1 against proteins identified in the HERD-1 IP-MS in this study. Dashed lines indicate the thresholds at 0.5 for the iPTM and 0.75 for the PEAKavg.

(B) AlphaFold3 prediction of HERD-1 (residues 1-718, green) with DEPS-1 (residues 1-619, yellow) including the PAE plots for all five models.

(C) Scheme of ZNFX-1 highlighting domain boundaries that were used for the HT-ColabFold screening shown in Figure 6A.

(D) AlphaFold3 prediction of DEPS-1 (residues 1-619, yellow) with ZNFX-1 (residues 1-190, blue) including the PAE plots for all five models.

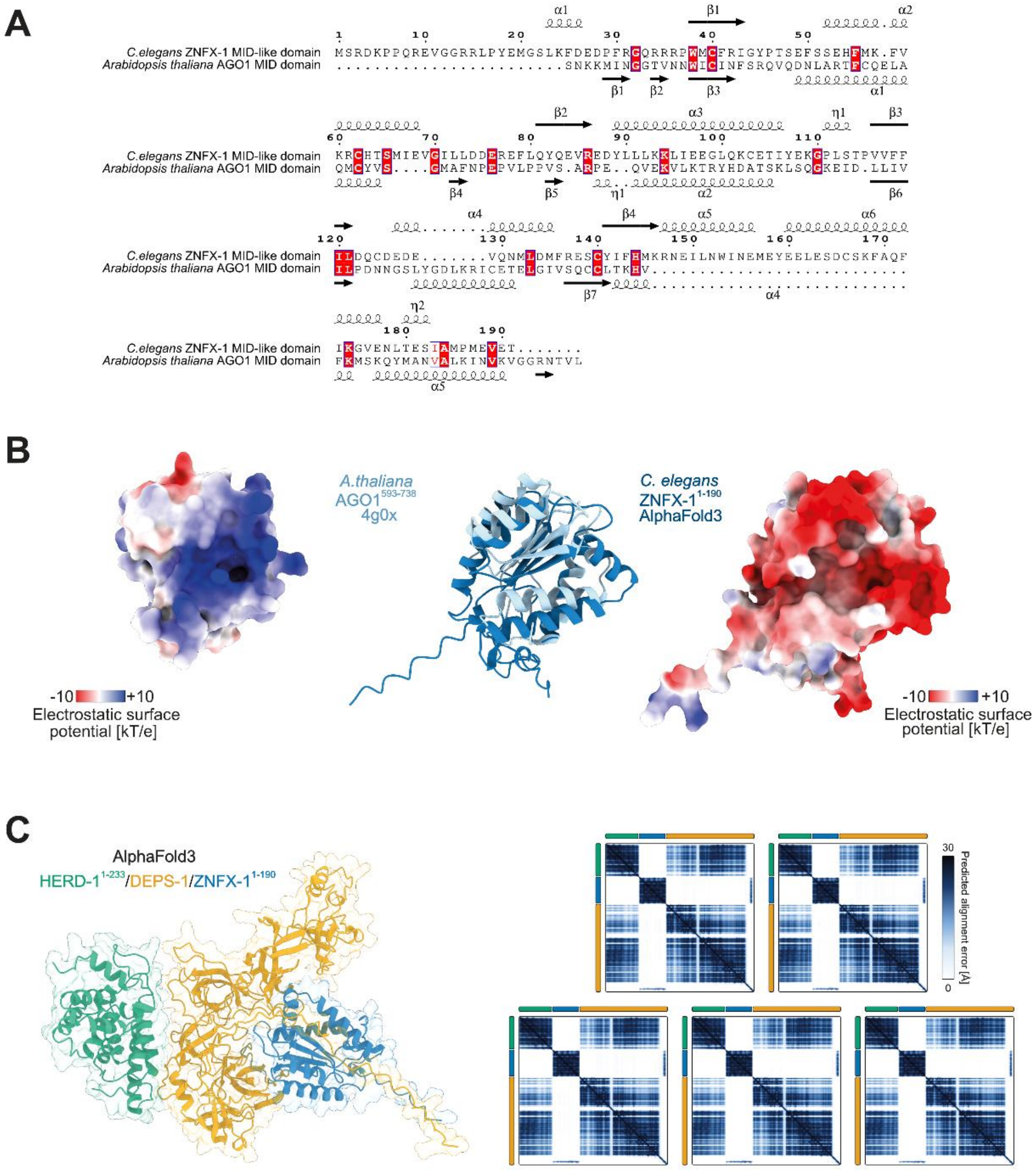


Pshanichnaya, L. *et al.*; Figure EV6

**Figure EV6. ZNFX-1 N-terminus has MID-like domain structure.**

(A) Sequence alignment of the MID-like domain of ZNFX-1 (residues 1-190) and the MID domain of AGO1 from *Arabidopsis thaliana* (residues 593-738). In the alignment, identical residues are highlighted in red boxes; residues with similar properties are indicated in white boxes. Helix represents α-helixes (α) and $3_{10}$-helices (η), and arrow – β-sheet. The alignment places the ZNFX-1 2D structure annotation above the sequence and the AGO1 annotation below it.

(B) Structural comparison between the MID-like domain of ZNFX-1 (residues 1-190, blue) predicted with AlphaFold3 and the crystal structure of the AGO1 MID domain from *Arabidopsis thaliana* (residues 593-738, PDB: 4g0x, light blue). RMSD between the model and the structure is 1.231 Å. Electrostatic surface potentials are a Coulomb summation over atomic partial charges carried out in ChimeraX.

(C) AlphaFold3 prediction of DEPS-1 (residues 1-619, yellow) and HERD-1 (residues 1-233, green) with ZNFX-1 (residues 1-190, blue) including the PAE plots for all five models.

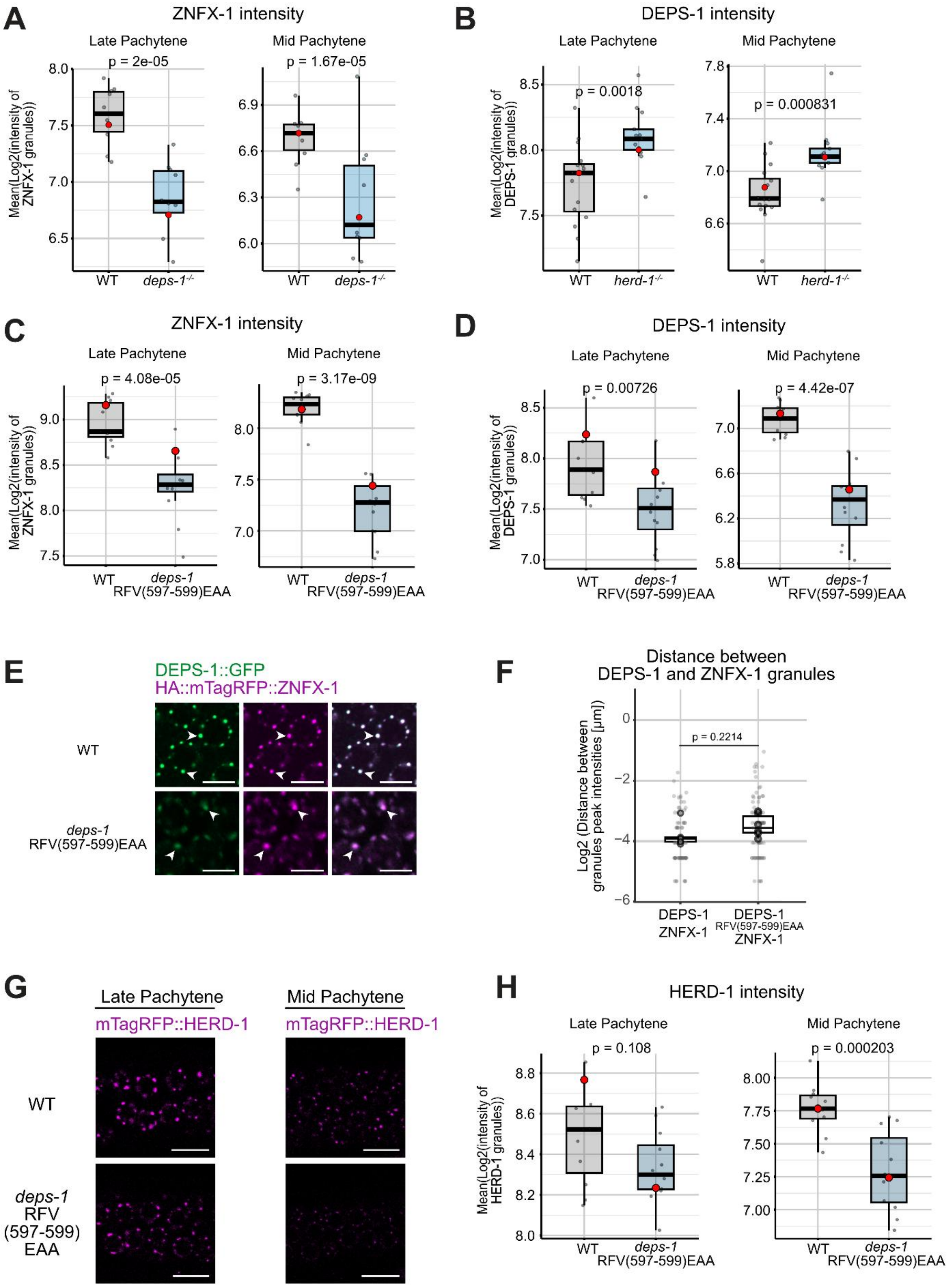


Pshanichnaya, L. *et al.*; Figure EV7

**Figure EV7. DEPS-1 is an important component of Z granules.**

(A-D,H) Quantification of ZNFX-1 (D), DEPS-1 (E) and HERD-1 (H) granule intensity shown in Figure 6C – (A), Figure 6C – (B), Figure 6C – (C,D) and Figure EV7G – (H), analyzed separately for the Mid and Late Pachytene regions. Each dot represents the mean log2-transformed granule intensity of an individual animal; boxes indicate the 25th–75th percentile range, and horizontal lines indicate the median, whiskers reach the most extreme non-outlier values. Red dots refer to representative micrographs shown in Figure 6, EV6G. Statistical significance was assessed using a linear mixed model. n = ~10 animals per condition.

(E) Representative fluorescence microscopy of the one nucleus in the pachytene germline region of young adult hermaphrodites expressing DEPS-1::GFP together with mTagRFP::ZNFX-1 in the indicated mutant backgrounds. Arrows indicate individual perinuclear germ granules. Images are representative of three independent biological replicates. Scale bars, 5 µm.

(F) Quantification of distances between peak intensities of granules shown in (E). Each light dot represents the distance between individual granules; dark dots represent the mean distance between granules within one worm; the box represents the 25th percentile and 75th percentile of the per-worm means, and horizontal bars indicate median values of the per-worm means, whiskers reach the most extreme non-outlier values. For logarithmic data representation, 0 values were replaced with half the pixel size value. Statistical significance was assessed using Welch's t-test. n = 5-6 biological replicates.

(G) Representative fluorescence microscopy of DEPS-1::GFP and mTagRFP::HA::HERD-1 in Mid and Late Pachytene regions of young adult animals in the indicated backgrounds. n = 11 biological replicates. Scale bars, 20 µm.

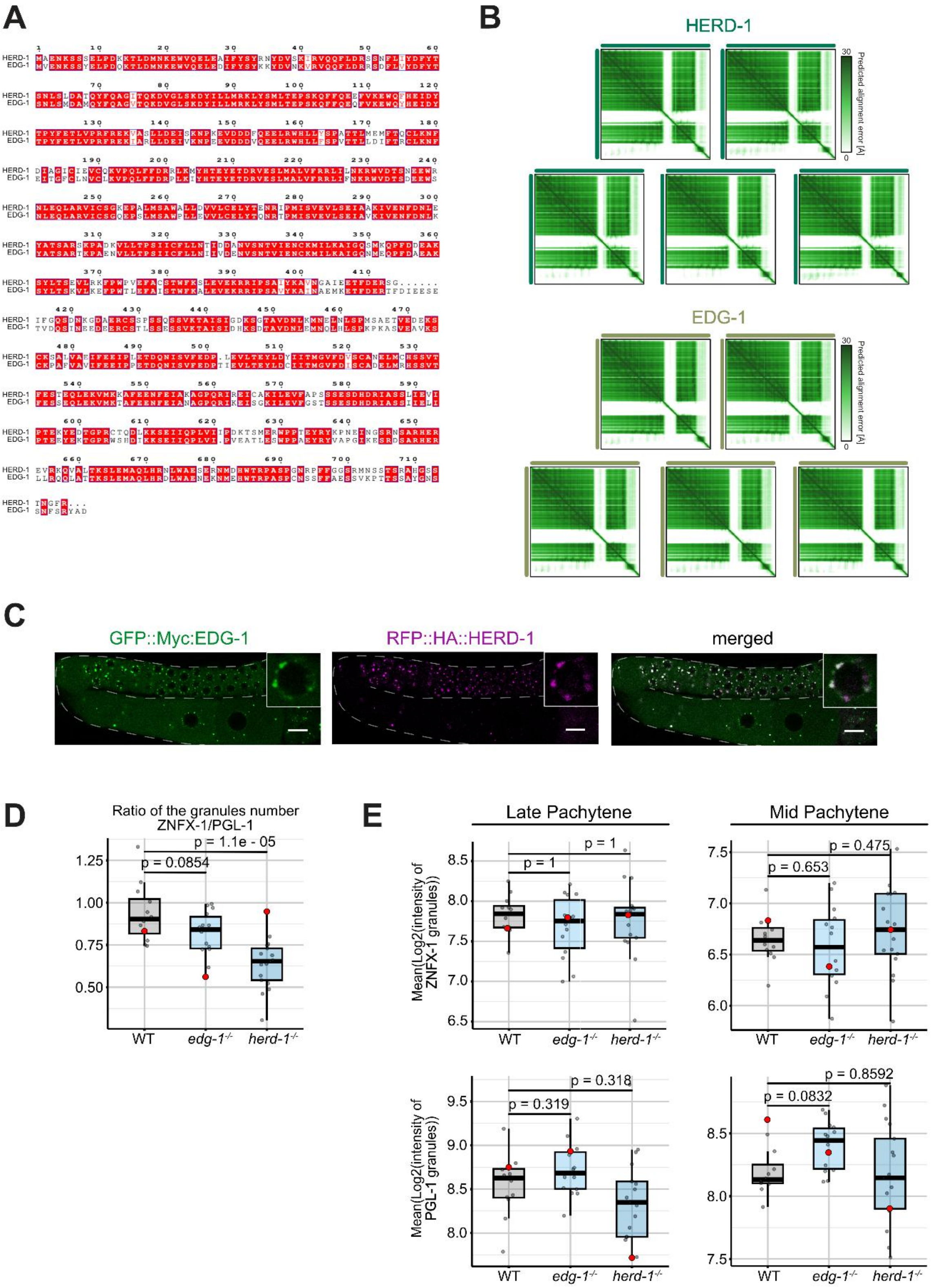


Pshanichnaya, L. *et al.*; Figure EV8

**Figure EV8.** ***edg-1****$^{-/-}$* **and** ***herd-1****$^{-/-}$* **single mutants do not affect granule intensity.**

(A) HERD-1 and EDG-1 protein sequence alignment. Identical residues are highlighted in red boxes; residues with similar properties are indicated in white boxes.

(B) Predicted Aligned Error (PAE) plots of AlphaFold3 structure from Figure 7A.

(C) Representative fluorescence microscopy of GFP::Myc::EDG-1 and RFP::HA::HERD-1 in the germline of young adult animals. Zoom-in shows germ granules around one nucleus. Dashed lines represent the germline boundary. Images are representative of three independent biological replicates. Scale bars, 10 μm.

(D) Quantification of granule number in the distal region of the germline. The number of ZNFX-1 granules was normalized to the number of PGL-1 granules. Each dot represents the granule ratio for an individual animal; boxes indicate the 25th–75th percentile range, and horizontal lines indicate the median. Statistical significance was assessed using Dunnett's multiple comparison test. n = 12–17 biological replicates.

(E) Quantification of ZNFX-1 and PGL-1 granule intensity shown in (Fig. 7D), analyzed separately for the mid and late pachytene regions. Each dot represents the mean log2-transformed granule intensity of an individual animal; boxes indicate the 25th–75th percentile range, and horizontal lines indicate the median, whiskers reach the most extreme non-outlier values. Red dots refer to representative micrographs shown in Figure 7D. Statistical significance was assessed using a linear mixed model, with Holm's correction for multiple comparisons. n = 12–17 animals per condition.
